# Context-Dependent Regulation of Microhomology-Mediated End Joining in Normal Tissues: Insights into Tissue-Specific Activation of DNA Repair Pathways

**DOI:** 10.1101/2025.05.23.653089

**Authors:** Diksha Rathore

## Abstract

Microhomology-mediated end joining (MMEJ) is a mutagenic DNA double-strand break (DSB) repair pathway, typically regarded as a backup mechanism in cancer, activated when canonical repair pathways such as non-homologous end joining (c-NHEJ) or homologous recombination (HR) are compromised. While MMEJ has been detected in normal tissues, its presence is puzzling given its error-prone nature, and its physiological role remains poorly defined. Recent studies implicating MMEJ in mitosis suggest a potential function in normal proliferative cells. Here, we show that MMEJ is not uniformly active across tissues but is selectively enriched in proliferative tissues, including thymus, spleen, testes, and liver, while markedly reduced in post-mitotic tissues such as brain, heart, kidney, and lung. This differential activity is supported by tissue-specific expression of key MMEJ components (e.g., Ligase III, MRE11, XRCC1, PARP1, Pol θ) and inhibitory factors (e.g., WRN, RAD51, ATM). Moreover, proliferative tissues preferentially utilize short microhomologies (∼10 nt), whereas post-mitotic tissues rely on longer microhomologies (≥13 nt), indicating a shift in repair pathway choice. These findings reveal that MMEJ is a tightly regulated, context-dependent repair pathway. Its activity is tolerated in proliferative tissues due to ongoing cell turnover, while its suppression in long-lived, post-mitotic cells is likely essential to preserve genomic stability. This study assigns a physiological role to MMEJ in healthy tissue homeostasis and highlights its relevance for designing targeted DNA repair-based therapeutic strategies across diverse tissue types.

## Introduction

Double-strand breaks (DSBs) are among the most detrimental forms of DNA damage, arising from endogenous and exogenous sources [1]. DSBs can be repaired through multiple pathways, including Homologous Recombination (HR), Non-Homologous End Joining (NHEJ), Microhomology-Mediated End Joining (MMEJ), and Single-Strand Annealing (SSA) [1][2]. While HR is highly accurate, NHEJ directly ligates DNA ends with minimal end processing, potentially leading to mutations [3][4]. MMEJ and SSA are more error-prone, frequently generating deletions and chromosomal translocations linked to cancer [5][6].

MMEJ begins with end resection by the MRN complex and CtIP, generating 3′ overhangs, and HMCES protects these overhangs [6][7][8]. Pol θ aligns microhomologous regions, displaces RPA, and stabilizes the annealed intermediate [9][10]. Recent studies indicate that additional polymerases, such as Pol λ and Pol β, also play roles in MMEJ. Pol θ and Pol δ cooperate during mitotic repair, with Pol δ aiding in gap filling [11][12][13]. Non-homologous flaps are removed by FEN1 and APE1 [14][15], followed by ligation through XRCC1–Ligase III, with Ligase I as a backup [16][17][18]. PARP1 enhances synapsis and promotes repair through poly(ADP-ribosyl)ation of repair proteins [19]. Additionally, p53, WRN, ATM, RAD51, RPA, and BRCA1 play regulatory roles [18][20][21][22].

MMEJ is a well-established contributor to genomic instability in cancer. Chromosomal translocations, such as VTI1A-TCF7L2 in colorectal cancer and Igh–c–Myc in lymphomas, often exhibit microhomology at breakpoints, reflecting MMEJ’s involvement [23][24]. Pol θ, a key MMEJ polymerase, is overexpressed in several cancers, including lung, colon, stomach, and glioblastoma, linking elevated MMEJ activity to tumor progression and poor prognosis [25][26][27].

While MMEJ is frequently associated with oncogenic processes, recent studies highlight its physiological function: MMEJ alleviates replication stress, repairs S-phase DSBs during mitosis [28][29][30], and contributes to antibody class-switch recombination [31][32][33]. In the absence of cNHEJ, MMEJ supports telomere maintenance and V(D)J recombination without RAG [34][35][36][37][38][39]. Its activity has been demonstrated in cell-free extracts from rat testes, calf thymus, and *Xenopus laevis* eggs [18][40][41]. Moreover, Pol θ expression in normal tissues such as testes, thymus, and placenta underscores the physiological relevance of MMEJ [42].

Genome-wide analyses reinforce this view, revealing that approximately 88% of protein-coding regions, comprising nearly 11 million deletions, are flanked by microhomologies [43]. The size of these deletions tends to increase with microhomology length, suggesting a shift in repair pathway usage [43]. MMEJ typically utilizes microhomologies ranging from 5 to 25 base pairs [44], though some studies report activity with sequences as short as 10 nucleotides [45][46]. Remarkably, the frequency of MMEJ events increases significantly—up to tenfold—for each additional base pair between 12 and 17 nucleotides [47]. In contrast, microhomologies longer than 30 nucleotides tend to promote single-strand annealing (SSA) [48]. However, the precise microhomology length threshold that mechanistically distinguishes MMEJ from SSA remains to be defined.

While isolated studies have reported MMEJ activity in normal tissues such as the thymus and testes [18], its prevalence and regulatory dynamics across a broader array of healthy tissues remain largely unexplored. The extent to which MMEJ is active in different tissues may vary significantly depending on the local repair environment and the unique cellular contexts of each tissue. For instance, tissues with high turnover, such as the thymus, may experience a higher frequency of DNA damage, potentially leading to increased reliance on DNA repair. In contrast, tissues with low replication rates may utilize these pathways less frequently. Moreover, the factors influencing MMEJ, including the length of microhomologies, might differ across tissues due to variations in DNA damage response signaling, the availability of repair proteins, and the local chromatin architecture.

Various proteins have been implicated in MMEJ through gene silencing approaches [18], but their roles in regulating MMEJ activity may vary depending on tissue-specific factors. For example, proteins involved in DNA resection or end processing might be differentially expressed or functionally modulated in different tissues. This leads to tissue-specific variations in MMEJ efficiency and repair events.

These observations highlight several unresolved aspects in the biology of MMEJ, including the microhomology threshold that defines MMEJ and distinguishes it from SSA. The extent to which MMEJ activity varies among different tissues remains an important area of investigation, particularly in understanding how this pathway contributes to genome maintenance in a tissue-specific manner and how it is regulated at the molecular level across diverse cellular contexts. Addressing these aspects is essential for understanding the tissue-specific roles of MMEJ in genome stability and repair pathway choice in mammalian cells.

The present study indicates tissue-specific differences in MMEJ. My data showed efficient and rapid repair in proliferative tissues such as the thymus, testes, spleen, and liver, compared to minimal activity and delayed repair in non-proliferative tissues like the brain, heart, lungs, and kidneys. This study also demonstrates that microhomology length and composition influence DNA repair efficiency across various tissues. Remarkably, the spleen preferentially utilizes 10 nt microhomology, whereas the brain, lungs, heart, and kidneys tend to favor more extended sequences. Furthermore, we characterize key factors—Ligase III, MRE11, XRCC1, FEN1, PARP1, and Pol θ—that regulate MMEJ and restrict its activity to dividing cells. These findings provide new insights into the tissue-specific regulation of MMEJ and its microhomology-length dependence, setting the stage for understanding how this repair pathway functions across different physiological contexts.

## Material and methods

### Enzymes, chemicals, and reagents

The chemicals and reagents used in the study were sourced from Amresco (USA), SRL (India), Sigma Chemical Co. (USA), and Himedia (India), and Restriction enzymes and other DNA-modifying enzymes were acquired from New England Biolabs (USA) and Fermentas (USA). Radioisotope-labelled nucleotides were supplied by BRIT (India) and Revvity (USA). Antibodies were procured from BD (USA), Cell Signaling Technology (USA), Santa Cruz Biotechnology (USA), and Calbiochem (USA). Oligomeric DNA was obtained from Eurofins Genomics (India), Juniper LifeSciences (India), Medauxin (India), and Xcelris Genomics (India).

### Oligomeric DNA

The oligomers used in this study are detailed in Table 1. These oligomeric DNA were purified using 12-15% denaturing PAGE whenever needed. The relevant section on materials and methods provides specific information regarding the use of oligomers in each experiment. For clarity, a simplified nomenclature is applied to some of these oligomers when presenting experimental results.

**Table 1:**
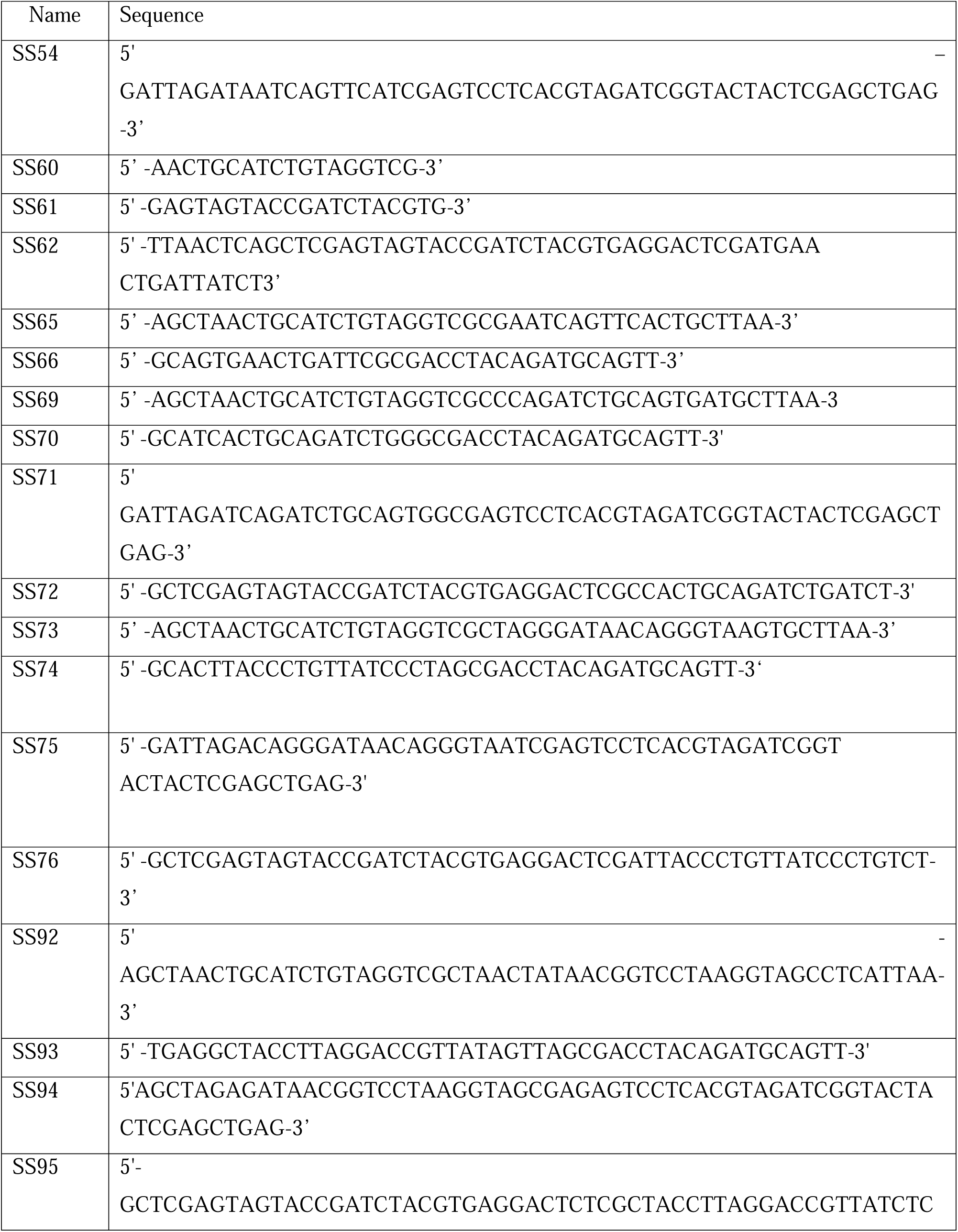
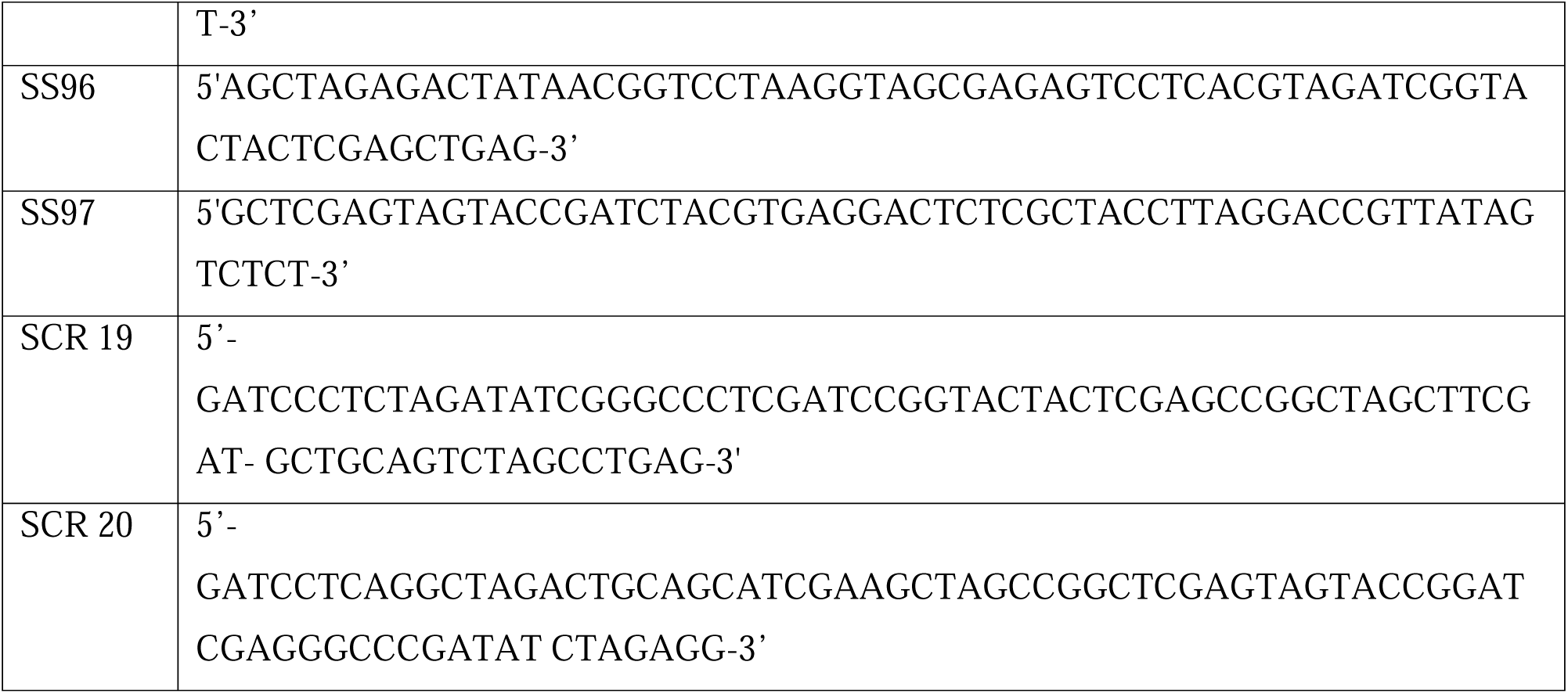

### Preparation of dsDNA substrates from oligomers

Oligomeric DNA mimicking double-strand breaks flanked by direct repeats was created by annealing complementary oligomers in a solution containing 100 mM NaCl and 1 mM EDTA, as previously described.^62^The study used microhomology regions of varying lengths: 10, 13, 16, 19, and 2 2 nucleotides (see Table 1). As described earlier, the 5′ end-labelling of SS60 and other oligomers with γ-32P ATP was performed using T4 polynucleotide kinase, and the labelled oligomers were stored at −20°C.

### Animals

Male Wistar rats (Rattus norvegicus) aged 4-8, 12–16, and 24-32 weeks were obtained from the Central Animal Facility at the Indian Institute of Science (IISc) in Bangalore, India. They were kept according to the Animal Ethics Committee guidelines (CAF/Ethics/ 871/2021) and complied with Indian national animal care and use regulations. The rats were housed in a controlled environment with regulated temperature and humidity and a 12 h light/dark cycle. They were kept in polypropylene cages and provided with a standard pellet diet (Agro Corporation, Pvt. Ltd., India) and water ad libitum. The pellet diet includes 55% nitrogen-free extract (carbohydrates), 21% protein, 5% lipids, 4% crude fibre, 8% ash, 1% calcium, 0.6% phosphorus, 3.4% glucose, and 2% vitamins.

### 5’ end labelling of oligomeric substrates

The 5’ end labelling of oligomeric substrates was done using γ-32P ATP in the presence of 1 U of T4 Polynucleotide kinase in a buffer containing 20 mM Tris-acetate [pH 7.9], 10 mM magnesium acetate, 50 mM potassium acetate, and 1 mM DTT at 37°C for 1 h. The radiolabelled oligomers were purified on a Sephadex G25 column and stored at -20°C [50, 73].

### Preparation of cell-free extracts from various rat organs

Cell-free extracts were prepared from the various rat organs, such as testes, brain, lungs, heart, spleen, kidneys, liver, and thymus of male Wistar rats aged 4–6 weeks, 12–16 weeks, and 24–32 weeks, as previously described [41]. Briefly, tissues were washed with PBS and then in hypotonic buffer (10 mM Tris⋅HCl, pH 8.0; 1 mM EDTA; 5 mM DTT) at 2000 rpm, 10 min, 4°C. They were resuspended in 2 volumes of hypotonic buffer and homogenized after 20 min. Protease inhibitors (phenylmethylsulfonyl fluoride at 0.01 M, aprotinin at 2 μg/ml, pepstatin at 1 μg/ml, and leupeptin at 1 μg/ml) were added, and the mixture was kept on ice for 20 min. Then, 0.5 volumes of high-salt buffer (50 mM Tris⋅HCl, pH 7.5; 1 M KCl; 2 mM EDTA; 2 mM DTT) were added. The extract was centrifuged for 3 h at 42,000 rpm at 4°C using a Beckman TLA-100 rotor. The supernatant was dialyzed overnight against dialysis buffer (20 mM Tris⋅HCl, pH 8.0; 0.1 M KOAc; 20% glycerol; 0.5 mM EDTA; 1 mM DTT), and the sample was snap-frozen and stored at −80°C.

Protein concentration was measured using Bradford’s assay and further normalized by loading on SDS-polyacrylamide gel, followed by staining with Coomassie Brilliant Blue.

### MMEJ assay

MMEJ reactions were performed as detailed previously [18, 50, 74]. Briefly, MMEJ substrate with various microhomology lengths (10, 13, 16, 19, and 22 nucleotides) was incubated with 0.5–3 µg of mitochondrial extracts or cell-free extracts in a buffer containing 50 mM Tris-HCl (pH 7.6), 20 mM MgCl2, 1 mM DTT, 1 mM ATP, and 10% PEG at 30°C/37°C or appropriate temperature and time. The reactions were ceased by heating at 65°C for 20 min to denature the proteins. The end-joined products were then PCR amplified using radiolabeled primer SS60 and unlabeled primer SS61 [denaturation: 95°C for 3 min (1 cycle); denaturation: 95°C for 30 s, annealing: 58°C for 30 s, extension: 72°C for 30 s (15 cycles); extension: 72°C for 3 min (1 cycle)]. The amplified joined products were analyzed using 10% denaturing PAGE. A 60-nt radiolabeled oligomer was included alongside the reactions as a marker, as the expected MMEJ product is 62 nt. The signals were detected with a phosphorImager (Fuji, Japan) and analyzed using Multi Gauge (V3.0) software.

### Cloning and sequencing of end-joined junctions

MMEJ reaction products were PCR amplified and resolved using 10% denaturing PAGE. Bands of interest were excised from the gel, and the DNA was purified using TE buffer and 5 M NaCl. Proteins were removed through a phenol: chloroform extraction, and the DNA was then precipitated. The purified DNA was ligated into a TA vector and incubated at 16°C for 16 h before being transformed into E. coli. Following transformation, plasmid DNA was isolated and digested to verify the presence of the insert, and positive clones were sequenced (Barcode Biosciences, India).

### Immunoblot analysis

For immunoblotting analysis, approximately 20–40 μg of protein was separated using 8–12% SDS-PAGE, as described by Chiruvella et al. (2008) [75]. After electrophoresis, the proteins were transferred to a PVDF membrane (Millipore, USA). The membrane was blocked with 5% non-fat milk or BSA in PBS containing 0.1% Tween-20. Proteins were then detected using specific primary antibodies against Cytochrome-C, PCNA, Tubulin, Lig3, Lig1, XRCC1, Rad50, Rad51, Polymerase theta, Parp1, FEN1, p53, MRE11, and others, followed by corresponding secondary antibodies according to standard protocols. The blots were developed with a chemiluminescent substrate (Clarity, Biorad) and visualized using a gel documentation system (LAS 3000, FUJI, Japan).

### Quantification

We used Multi Gauge (V3.0) software to measure the bands of interest. We selected a rectangle around each band to measure its intensity and did the same for all bands in each lane. We also measured the background intensity from a blank area of the same lane and subtracted it. The final intensity values were plotted and shown in a bar graph.

### Statistical analyses

Data from experiments carried out with a minimum of three repeats were subjected to statistical analysis by either Student’s t-test or analysis of variance (ANOVA), followed by Tukey or Dunnett’s post hoc test using GraphPad Prism 6 software (San Diego, CA, USA).] Results were considered statistically significant if the p-value was ≤0.05.

## Results

### MMEJ was most pronounced in the thymus, spleen, and testes, with lower activity observed in the lungs and kidneys using a 10 nt microhomology substrate

To investigate the presence and variability of MMEJ repair across different rat tissues, we prepared cell-free extracts from various tissues, including the testes, brain, lungs, heart, spleen, kidneys, liver, and thymus. Cell-free extracts were prepared as described previously [49] and normalized using 8% SDS-PAGE for subsequent analyses (Figure 2a and 2b). In vitro MMEJ assays were conducted as outlined by Sharma et al. (2015) [18]. Briefly, two distinct oligomeric DNA substrates (SS54/62 and SS65/66), each containing a 10 nt microhomology region designed to mimic a DSB flanked by direct repeats, were incubated with normalized CFE for 1 h at 37°C (the physiological temperature for most tissues). Incubation for testes extracts was performed at 30°C (a lower temperature that better supports spermatogenesis) (Figure 1a). Microhomology within the substrate directs repair via MMEJ, generating shorter products due to deletion at the joining site (Figure 1b). The resulting joined products can be detected by radioactive PCR. A 62-nt joining product was observed in the presence of CFE (Figure 2c).

**Fig. 1.**
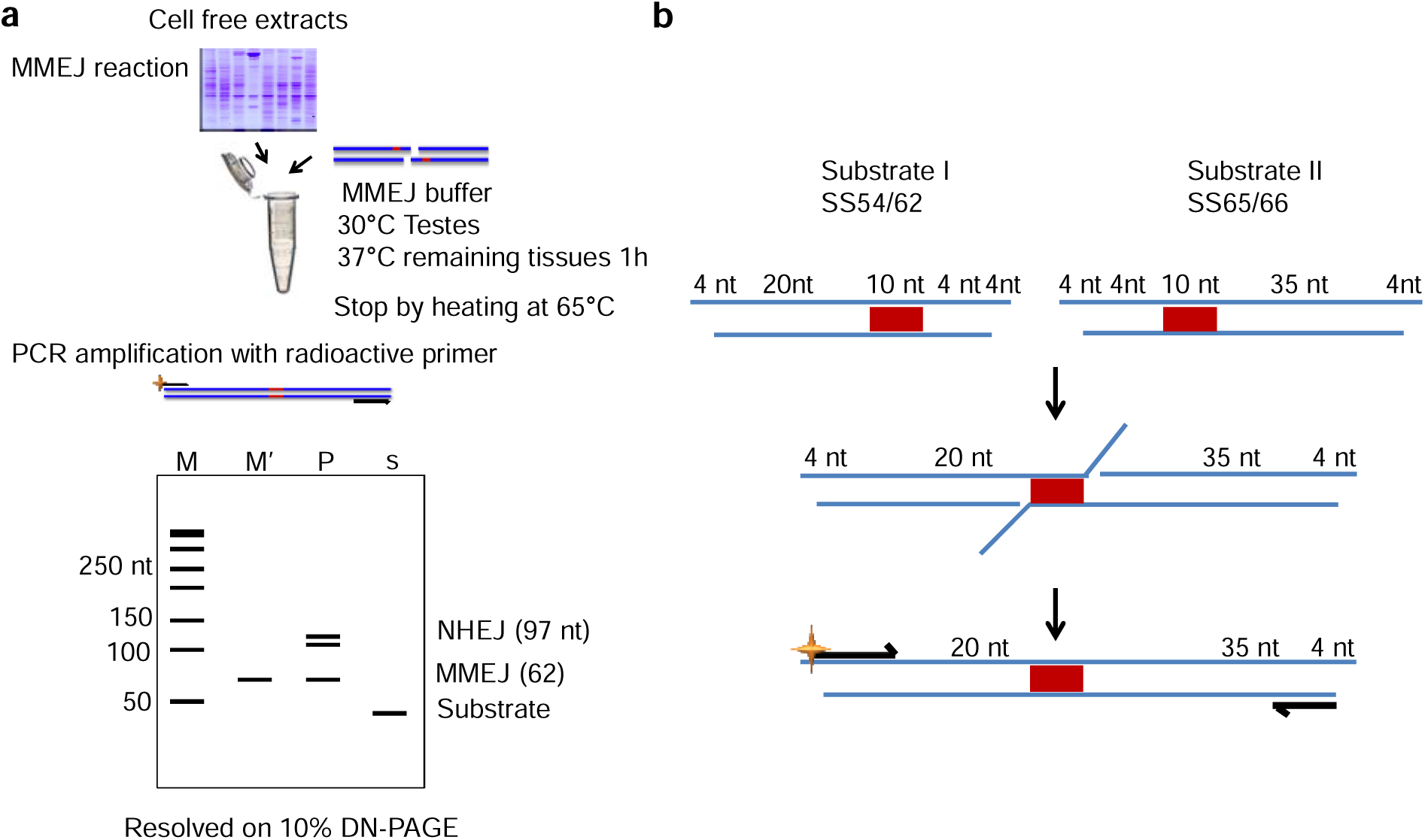
Schematic representation of the *in vitro* Microhomology-Mediated End Joining (MMEJ) assay. (a) The diagram outlines the steps involved in the MMEJ assay. Two substrate DNA molecules, SS54/62 and SS65/66, were incubated with normalized cell-free extract (CFE) from various rat organs in buffer at the appropriate temperature for 1 h. The reaction was terminated by heat inactivation, and the join product was amplified using radioactive primers. The resulting PCR products were resolved on a 10% denaturing PAGE. M represents the 50 nt marker, M’ is the 60 nt marker, ‘S’ indicates the substrate, and ‘P’ means the product formed due to repair. (b) The schematic representation of the substrate used in the assay. These substrates are designed to mimic a double-strand break, with 10-nt microhomology flanking the break site. Following joining via MMEJ, the final product is reduced to approximately 62 nt due to the deletion of one microhomology and the adjacent sequence, resulting in a product containing a single microhomology at the break site. The red box highlights the microhomology region.

**Fig. 2.**
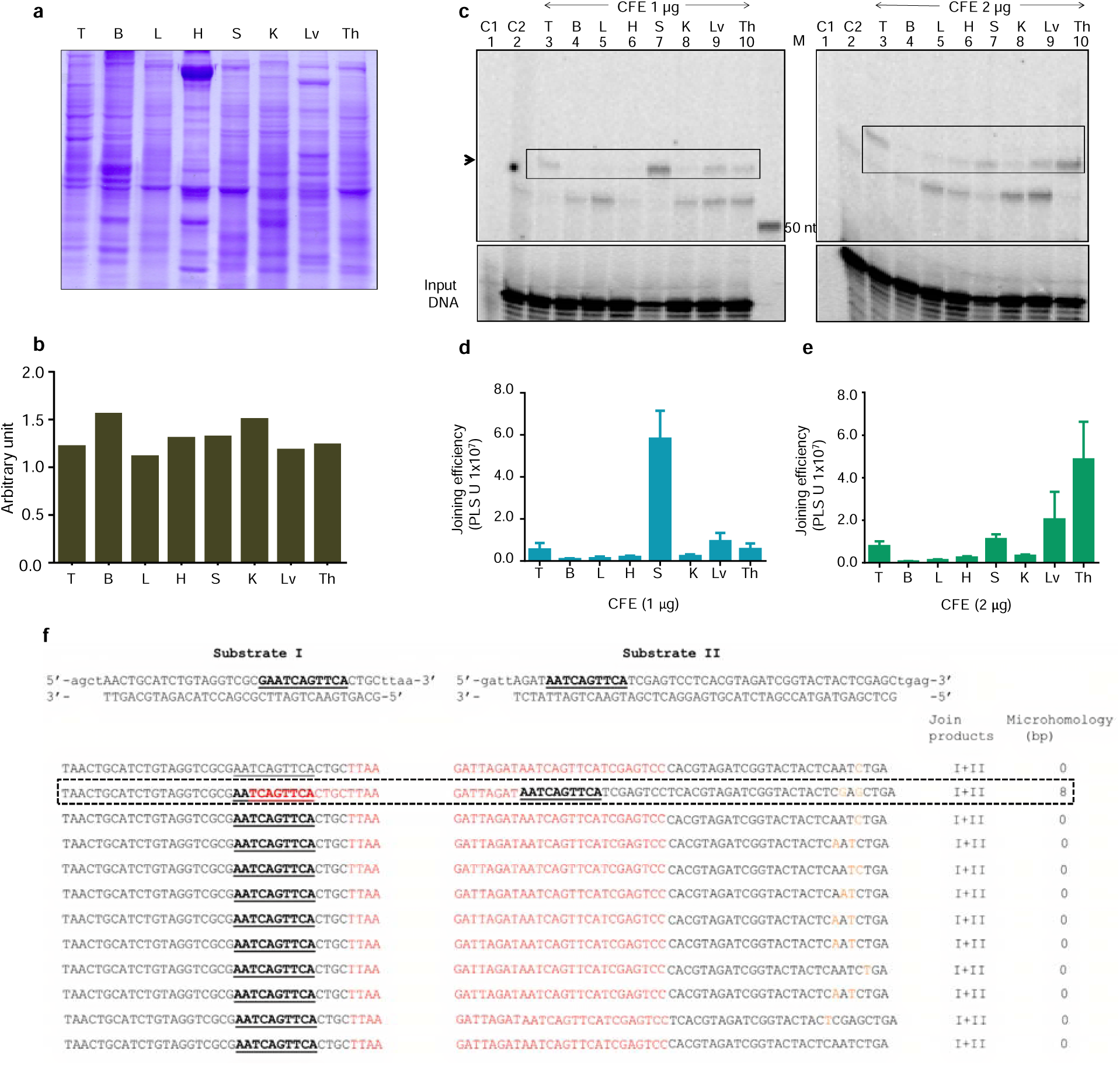
Comparison of the joining efficiencies of different rat organs: SDS-PAGE profiles were generated using cell-free extracts (CFE) prepared from various rat tissues, including testis (T), brain (B), lungs (L), heart (H), spleen (S), kidneys (K), liver (Lv), and thymus (Th) (panel a). A bar graph showing the normalization profile for protein content based on SDS-PAGE is presented in panel (b). Panel (c) illustrates the MMEJ efficiency comparison among different organ CFEs using 1 µg and 2 µg of extract incubated with a substrate mimicking a double-strand break (DSB) flanked by 10-nucleotide direct repeats, followed by analysis on 10% denaturing PAGE with a 50-nt ladder (M). Panels (d) and (e) show bar graphs representing MMEJ efficiencies catalyzed by 1 µg and 2 µg CFE, respectively, from each organ, relative to input DNA. In both cases, a minimum of three biological replicates were performed, and error bars indicate the mean ± SEM. In panel (f), the join products corresponding to MMEJ were excised from the gel, DNA was extracted, PCR amplified, cloned, and sequenced. Each sequence shown is derived from an independent clone. Instances of microhomology utilization are highlighted with a dotted box, where the microhomology sequences are bold, black, and underlined; deleted regions are shown in red, and mutations are marked in orange. A total of 12 clones were sequenced.

No joining was detected in reactions lacking protein (Lane 2, Figure 2c), thereby confirming that the observed joining is catalyzed by the protein within the CFE (Figure 2c). Quantification of joint products revealed tissue-specific variations in repair efficiency. When 1 µg of CFE was used, the highest level of joining was observed in the spleen, followed by the liver, thymus, and testes (Figure 2c and 2d). In contrast, when 2 µg of CFE was used, the thymus exhibited the maximum joining, followed by the liver, spleen, and testes (Figure 2c & e). Interestingly, minimal or no joining was detected in the brain, lungs, heart, and kidneys at 1 µg and 2 µg CFE concentrations (Figure 2c, lanes 4, 5, 6, and 8, respectively). These results suggest that DNA repair efficiency, as measured by MMEJ, may not be a uniform process but might instead be tissue-specific and potentially influenced by differences in cellular composition.

The higher repair activity was observed in tissues predominantly containing proliferative cells, such as the thymus, spleen, testes, and liver, and minimal repair in tissues composed mainly of non-dividing or post-mitotic cells, including the brain, lungs, heart, and kidneys.

To exclude the possibility that reduced end-joining activity in post-mitotic tissues was due to compromised CFE function, we conducted a control NHEJ assay using CFEs prepared from various rat organs. The activity profile of these extracts was consistent with earlier findings ^18, 73^, confirming their functional integrity (Supplementary Figure 1 b and d).

This study was conducted using three independent batches of CFE from each organ. To account for potential variation in DNA repair efficiency between batches, we normalized the CFE from each organ across all three batches. The normalized CFE was then used to substantiate the differences in DNA repair observed in Figure 2. When equal amounts of CFE from each organ were used across individual batches, we found that joining efficiency was consistent between batches, except the spleen, where one batch exhibited stronger joining (Supplementary Figure 2a).

### Minor repair junctions showed canonical MMEJ features, while atypical deletions suggest involvement of non-canonical MMEJ processes

To confirm whether the joined products observed in Figure 2 were generated via MMEJ, we excised the corresponding gel bands, purified the DNA, and cloned the products into a TA vector for sequencing. Junction analysis from three independent biological replicates revealed canonical MMEJ signatures, including the deletion of flanking DNA sequences between two direct repeats, resulting in a final product that retains a single direct repeat at the junction [18, 50, 51]. Unexpectedly, many junctions exhibited extensive deletions that cNHEJ or HR could not explain. NHEJ typically results in minimal sequence loss and does not produce extensive Deletions were observed in the repair products, consistent with MMEJ activity [52]. While homologous recombination (HR) typically requires long homology tracts (∼70 nt), which were absent from our substrate design [53], the deletions observed in our study exceeded the typical range associated with known MMEJ activity. This raises the possibility that an alternative or less well-characterized repair mechanism may also contribute. Interestingly, only a minority of junctions exhibited the canonical MMEJ features—deletion of flanking sequences and retention of a single microhomologous repeat at the junction. This observation suggests that while MMEJ is active in normal tissues, its engagement is likely tightly regulated, possibly as a cellular strategy to limit mutagenic repair and preserve genomic integrity.

### Testes, spleen, and thymus exhibit MMEJ repair at low protein concentrations, while brain, lungs, and kidneys show minimal or undetectable activity

The differences in maximum joining efficiency between 1 µg and 2 µg of CFE (Figure 2 c,d & e) pointed to tissue-specific variations in MMEJ repair activity, suggesting that each organ may have a distinct optimal CFE concentration for maximal repair. We incubated a 10-nt homology substrate with increasing amounts of CFE (0.1-3 µg) from each organ to identify these optimal concentrations. Notably, joined products first appeared at just 0.1 µg of CFE in the testes, with peak activity between 0.5–1 µg (Figure 3a). However, maximal repair efficiency was observed in the thymus, spleen, liver, and heart at 2–3 µg of CFE (Figures 3c & 3d). Interestingly, minimal MMEJ activity was observed in the thymus and heart at 2 µg of CFE (Figure 3c, lane 6; Figure 3d, lane 7). In contrast, MMEJ was detectable at 0.5 µg in the spleen (Figure 3c, lane 10) and at 1 µg in the liver (Figure 3d, lane 12).

**Fig. 3.**
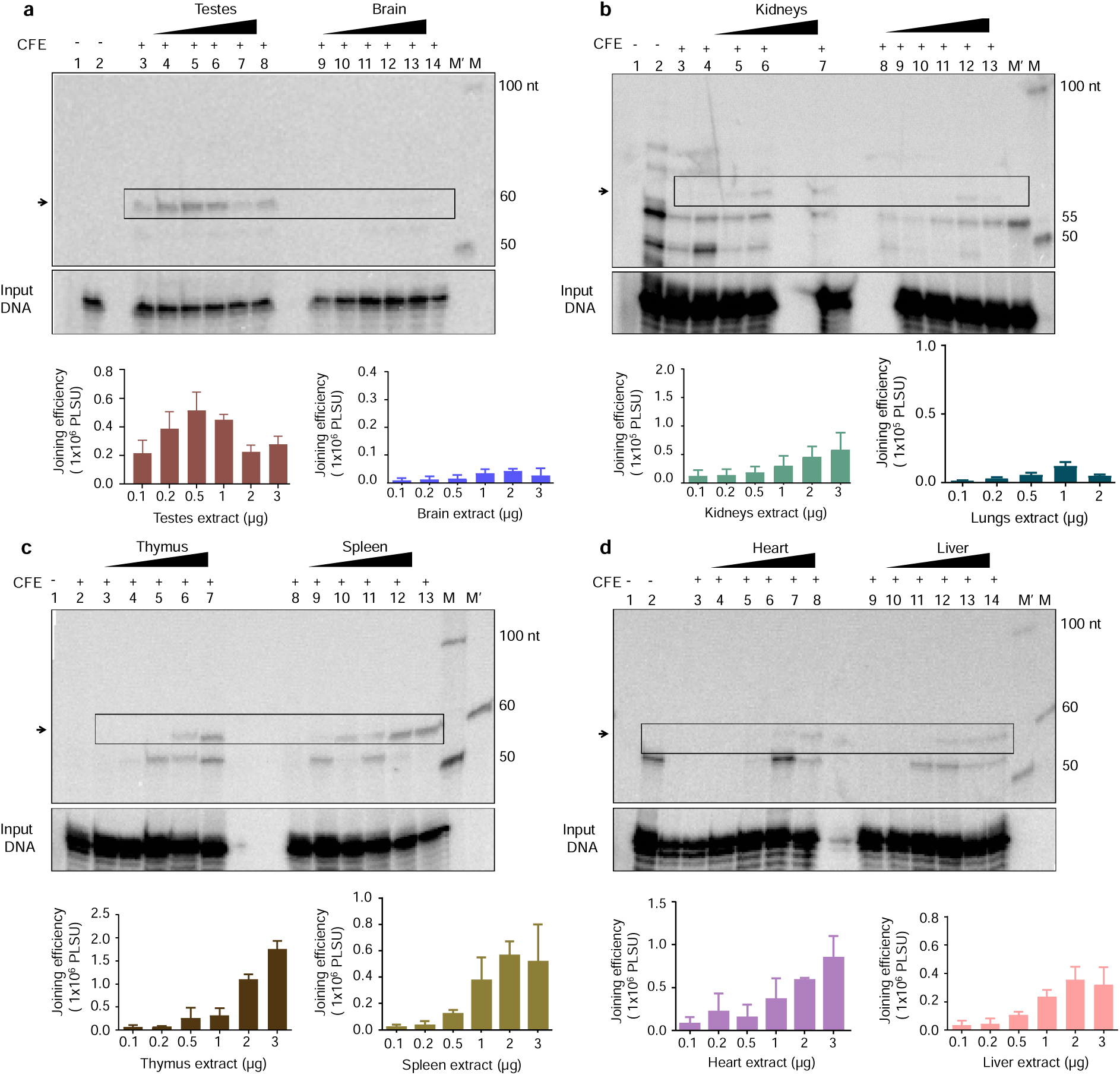
Evaluation of MMEJ activity in various rat tissue extracts at increasing protein concentrations using 10-nt microhomology DNA substrates. **(a)** Evaluation of MMEJ efficiency using increasing concentrations of cell-free extracts. Rat testicular (lanes 3-8) and brain extracts (lanes 9-14) (0.1, 0.2, 0.5, 1.0, 2.0, and 3.0 µg) were incubated with DNA substrates (4 nM) containing 10 nt microhomology for 1 h at 30°C (testes) and brain 37°C. Lane 1 represents the no-template control; lane 2 is the no-protein control. (**b**) Evaluation of MMEJ efficiency using increasing concentrations of cell-free extracts. Rat kidney (lanes 3-7) (0.1, 0.2, 0.5, 1.0, 2.0 and 3.0 µg) and lung extracts (lanes 8-13) (0.1, 0.2, 0.5, 1.0, and 2.0 µg) were incubated with DNA substrates (4 nM) containing 10 nt microhomology for 1 h at 37°C. Lane 1 represents the no-template control; lane 2 is the no-protein control. (**C**) Evaluation of MMEJ efficiency using increasing concentrations of cell-free extracts. Rat thymus (lanes 2-7) and spleen extracts (lanes 8-13) (0, 0.1, 0.2, 0.5, 1.0, 2.0, and 3.0 μg) were incubated with DNA substrates (4 nM) containing 10 nt microhomology for 1 h at 37°C. Lane 1 represents the no-template control. (**d**) Evaluation of MMEJ efficiency using increasing concentrations of cell-free extracts. Rat heart (lanes 3-8) and liver extracts (lanes 9-14) (0, 0.1, 0.2, 0.5, 1.0, 2.0 μg and 3.0 μg) were incubated with DNA substrates (4 nM) containing 10 nt microhomology for 1 h at 37°C. Lane 1 represents the no-template control; lane 2 is the no-protein control. Panel **(a-d)** A bar graph showing quantification from at least three independent biological experiments. An arrow and a box mark MMEJ products. The lower panel shows the loading control for equal DNA in each sample, labeled input DNA. M’ and M represent the 60 nt marker corresponding to the MMEJ join product and 50 nt ladder, respectively. The axis of the bar graph is labeled in photostimulated luminescence units (PSLU). Error bars represent the standard error of the mean (SEM).

In striking contrast, the lungs and kidneys exhibited minimal end-joining activity at 2 µg of CFE, with no substantial increase at higher concentrations (3 µg) (Figure 3b). No joining was observed in the brain, even at 3 µg of CFE (Figure 3a). These results highlight the distinct tissue-specific protein requirements for optimal MMEJ. The testes and spleen demonstrated efficient repair even at lower protein concentrations, while the thymus required comparatively higher CFE concentrations to achieve effective MMEJ. In contrast, the brain, lungs, and kidneys exhibited either very low or undetectable repair activity, regardless of increased CFE concentrations. These findings reinforce the earlier observation (Figure 2) that MMEJ activity varies across tissues and suggest that in specific organs, particularly those composed of non-dividing cells, MMEJ may be intrinsically constrained or tightly regulated, potentially as a protective mechanism to limit mutagenic repair outcomes.

### Tissue-specific variation in DNA repairs, with rapid initiation in high-MMEJ organs (thymus, spleen, and testes) and delayed repair in low-MMEJ tissues (lungs, kidneys)

Tissues such as the heart, lungs, kidneys, and brain exhibited minimal or no joining, even at the highest concentrations of CFE (Figure 3). This observation prompted further investigation into whether extending the incubation time could enhance repair efficiency in organs with low repair efficiency. To assess the time-dependent repair efficiency, we maintained a constant CFE concentration of 2 µg for each organ (which had previously shown optimal repair in most tissues) and a standardized incubation temperature of 37°C for all tissues, except the testes, which were incubated at 30°C to align with their physiological temperature. A 10-nt microhomology substrate was incubated with CFE from different organs for a range of durations: 3.75, 7.5, 15, 30 mins, 1 h, 2 h, and 4 h (Figure 4).

**Fig. 4.**
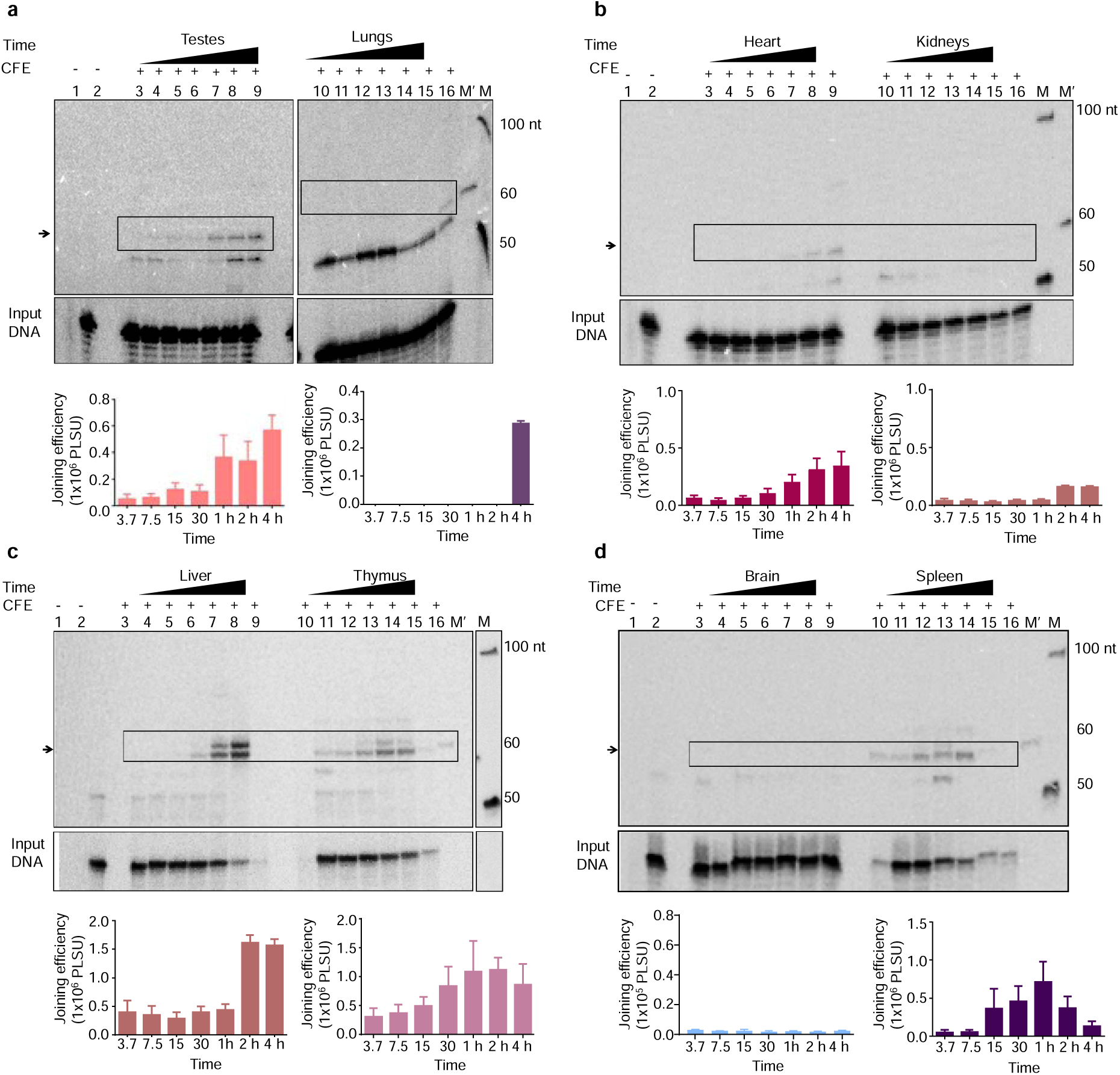
Time Kinetics of MMEJ on 10-nt Microhomology DNA Substrates in various rat tissue extracts: Time course analysis of MMEJ on DNA substrates containing 10-nt microhomology. Rat testicular (lanes 3-9) and lung extracts (lanes 10-16) (2 μg) were incubated with DNA substrates for 3.75, 7.5, 15, and 30 mins, as well as 1, 2, and 4 h, followed by an analysis of the products on 10% denaturing PAGE. **(b)** Time course analysis of MMEJ on DNA substrates containing 10-nt microhomology. Rat heart (lanes 3-9) and kidney extracts (lanes 10-16) (2 μg) were incubated with DNA substrates for 3.75, 7.5, 15, and 30 mins, as well as 1, 2, and 4 h, followed by an analysis of the products on 10% denaturing PAGE. **(c)** Time course analysis of MMEJ on DNA substrates containing 10-nt microhomology. Rat liver (lanes 3-9) and thymus extracts (lanes 10-16) (2 μg) were incubated with DNA substrates for 3.75, 7.5, 15, and 30 mins, as well as 1, 2, and 4 h, followed by product analysis on 10% denaturing PAGE. **(d)** Time course analysis of MMEJ on DNA substrates containing 10-nt microhomology. Rat brain (lanes 3-9) and spleen extracts (lanes 10-16) (2 μg) were incubated with DNA substrates for 3.75, 7.5, 15, and 30 mins, as well as 1, 2, and 4 h, followed by product analysis on 10% denaturing PAGE. Panel (**a-d**) bar graphs showing quantification from at least three independent experiments are presented. An arrow and a box mark MMEJ products. The lower panel shows the loading control for equal DNA in each sample, labeled ‘input DNA.’ M’ and M represent the 60 nt marker corresponding to the MMEJ join product and 50 nt ladder, respectively. The axis of the bar graph is labeled in photostimulated luminescence units (PSLU). Error bars represent the standard error of the mean (SEM).

Interestingly, early repair activity was detected in the testes, thymus, and spleen, with detectable joining starting at 3–7 minutes. In the testes, repair activity begins early and continues to rise throughout the incubation period (Figure 4a). In contrast, the thymus and spleen demonstrated an early initiation of repair, detectable as early as 3 mins, which increased with time and subsequently declined (Figures 4c & 4d). The heart and liver showed delayed repair initiation, detectable only after 1 h, with repair activity gradually increasing thereafter (Figures 4b & 4c). The kidneys showed minimal joining, even after 4 h of incubation, while the lungs demonstrated detectable repair only at 4h (Figures 4a & 4b). No joining was observed in brain extracts, even after 4 h of incubation, indicating a lack of MMEJ activity (Figure 4d).

These findings extend my earlier observations of tissue-specific differences in MMEJ efficiency (Figure2) and suggest that both the extent and timing of MMEJ may vary across tissues. Tissues like the testes, thymus, and spleen showed relatively rapid repair initiation. In comparison, the heart and liver exhibited a slower but sustained increase in repair activity over time. Meanwhile, kidneys and lungs showed limited repair, with detectable activity emerging only after extended incubation. These results suggest that MMEJ activity may be influenced by tissue-specific factors, including cellular turnover rates, metabolic demands, or the need to regulate DNA repair in specific physiological contexts tightly.

### Most organs exhibit the highest MMEJ repair at 25-30°C

Temperature is a critical extrinsic factor regulating DNA repair, with repair enzymes, including DNA ligases, demonstrating efficient ligation activity even at sub-physiological temperatures, such as 16°C [54]. In contrast, these enzymes function in vivo at 37°C, raising the possibility that temperature may modulate repair efficiency in a tissue-specific manner. Given the potential influence of local microenvironmental conditions, we hypothesized that the efficiency of MMEJ is both tissue and temperature-dependent. To investigate this, we performed a temperature titration, maintaining a constant protein concentration (2 µg CFE) in each organ and incubation time (1 h) across the experiment, as these conditions typically yielded optimal activity for most tissues. The incubation temperature was set from 4°C, 16°C, 25°C, 30°C, 37°C, and 42°C, with 37°C representing the physiological temperature for most tissues, except the testes, which function optimally at 30°C. MMEJ activity was assessed in different tissues at these temperatures (Figure 5a-d).

**Fig. 5.**
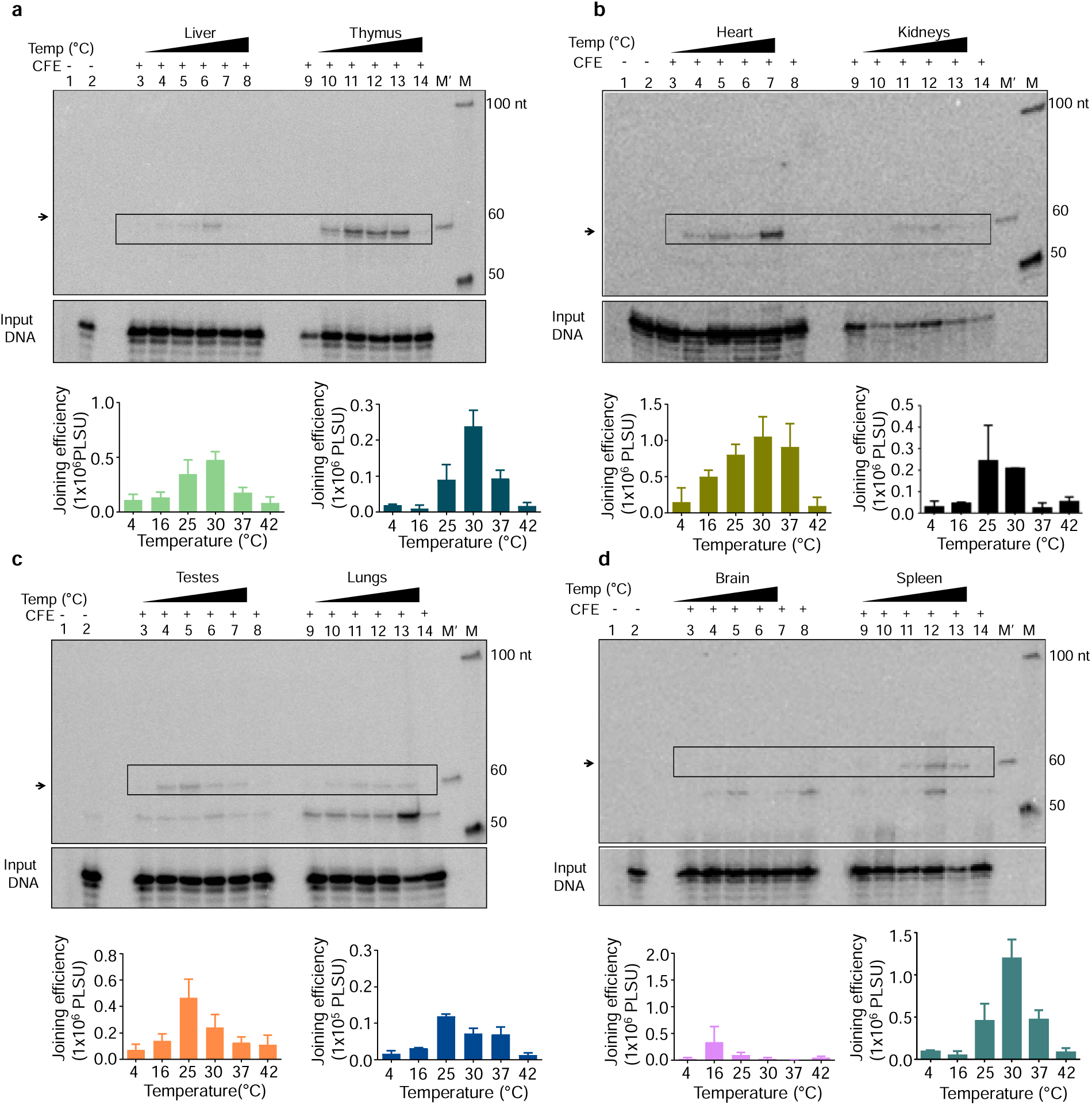
MMEJ assays at various incubation temperatures using different rat tissue extracts with 10-nt microhomology-containing DNA substrates. **(a)** MMEJ assay at increasing incubation temperatures. A 2.0 μg of liver and thymus extracts were incubated with 10-nt microhomology-containing DNA substrates for 1 h at 4, 16, 25, 30, 37, and 42°C. Lane 1 represents the no-template, and lane 2 represents the no-protein control. **(b)** MMEJ assay at increasing incubation temperatures. A 2.0 μg of heart and kidney extracts were incubated with 10-nt microhomology-containing DNA substrates for 1 h at 4, 16, 25, 30, 37, and 42°C. Lane 1 represents the no-template, and lane 2 represents the no-protein control. **(c)** MMEJ assay at increasing incubation temperatures. A 2.0 μg of testes and lung extracts were incubated with 10-nt microhomology-containing DNA substrates for 1 h at 4, 16, 25, 30, 37, and 42°C. Lane 1 represents the no-template, and lane 2 represents the no-protein control. **(d)** MMEJ assay at increasing incubation temperatures. 2.0 μg of brain and spleen extracts were incubated with 10-nt microhomology-containing DNA substrates for 1 h at 4, 16, 25, 30, 37, and 42°C. Lane 1 represents the no-template, and lane 2 represents the no-protein control. Panel (**a-d)** bar graphs showing quantification from at least three independent experiments are presented. An arrow and a box mark MMEJ products. The lower panel shows the loading control for equal DNA in each sample, labeled ‘input DNA.’ M’ and M represent the 60 nt marker corresponding to the MMEJ join product and 50 nt ladder, respectively. The axis of the bar graph is labeled in photostimulated luminescence units (PLSU). Error bars represent the standard error of the mean (SEM).

The results indicated that the liver, kidneys, and spleen exhibited temperature-dependent joining, with peak repair at 30°C. Joining efficiency decreased at lower and higher temperatures (Figure 5a, b, & d). In contrast, the thymus, heart, testes, and lungs demonstrated relative temperature resilience, maintaining MMEJ activity across a broader thermal range. For example, the thymus, heart, testes, and lungs showed peak repair at 25–37°C (Figure 5a, 5b & 5c). Intriguingly, these organs also maintained detectable MMEJ at 16°C with activity persisting up to 37°C (Figure 5a, b &c). This suggests that MMEJ in brain tissue may be highly restricted or selectively active under narrow, low-temperature conditions. Interestingly, although 37°C is considered physiological temperature, most tissues showed maximum *in vitro* MMEJ activity at lower temperatures, particularly 25–30°C. This likely reflects differences between *in vitro* and *in vivo* conditions. *In vitro* assays lack cellular compartmentalization, molecular crowding, and regulatory pathways. Moreover, lower temperatures may suppress nuclease activity, leading to greater substrate stability and improved access for repair enzymes, thereby enhancing ligation efficiency.

These results show that DNA repair via MMEJ is temperature-sensitive and resilient, with tissue-specific temperature preferences. Tissues like the liver, kidneys, and spleen peak at 30°C, while others, such as the thymus, heart, testes, and lungs, maintain activity across a broader range, including lower temperatures. While these findings are based on *in vitro* assays and do not necessarily reflect *in vivo* repair dynamics, they offer valuable insight into the biochemical behavior of MMEJ activity in different tissue extracts. The observed differences may be influenced by each tissue’s molecular composition and its response to temperature changes under simplified assay conditions.

### MMEJ activity is influenced by microhomology length and distinct tissue-specific responses observed across different organs

Although MMEJ can utilize microhomologies as short as 6 bp, its efficiency increases significantly with longer sequences, particularly between 12 and 17 nt [47]. While MMEJ typically operates within a range of 5–25 bp, SSA requires microhomologies longer than 30 nt. Nevertheless, some studies suggest that MMEJ can also function effectively with microhomologies as short as 10 nt [45–48]. This conflicting evidence has made it challenging to fully understand the role of microhomology length in repair pathway selection. To address this uncertainty, we investigated the impact of microhomology length on MMEJ activity using DSB-mimicking substrates with defined microhomologies of 10, 13, 16, and 19 nt (Figure 6). These substrates were incubated with 2 µg of CFE from various tissues, generating repair products of 62, 65, 68, and 71 nt in length, respectively. This approach allowed us to systematically examine the effect of microhomology length on end-joining efficiency and to investigate whether tissues exhibit distinct preferences for specific microhomology lengths.

**Fig. 6.**
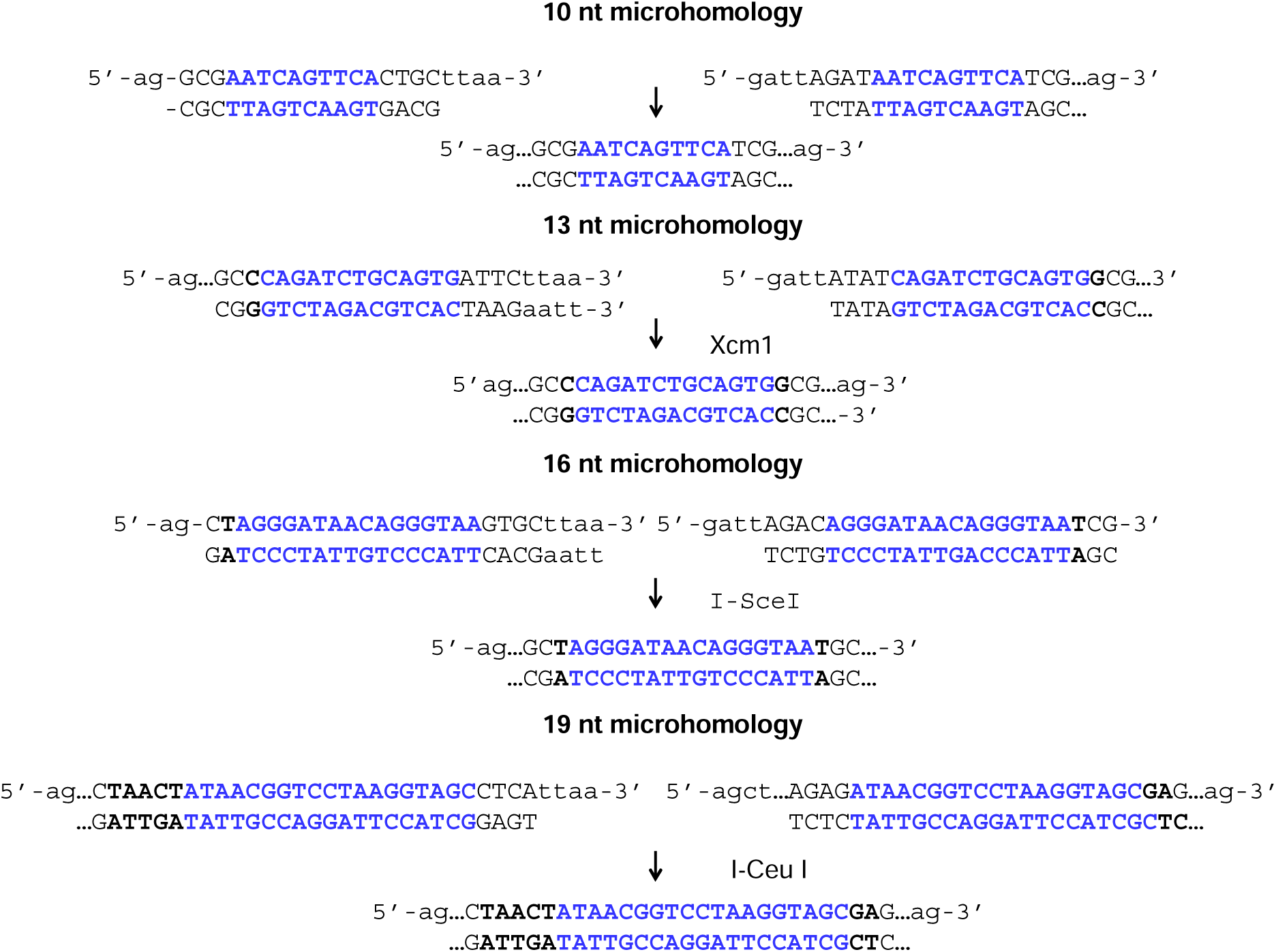
Schematic representation of DNA substrates with increasing microhomology length. DNA substrates were designed with microhomology regions of 10, 13, 16, and 19 nucleotides. Each substrate contains identical sequences upstream and downstream of the microhomology (shown in black), while only the length of the microhomology region (shown in bold blue) varies. A restriction site embedded within the microhomology enables verification that the repair products result from microhomology-mediated end joining. The schematic also shows the expected joined products formed through MMEJ.

Perhaps this study’s most striking and unexpected finding is that microhomology length acts as a key and tissue-specific regulator of MMEJ efficiency, revealing a level of complexity in DNA double-strand break repair that was previously unappreciated. My data demonstrate that its activity varies dramatically depending on the tissue type and the microhomology length (Figure 7). Using the substrates described above, we observed significant, reproducible shifts in repair efficiency across different tissues. The thymus, spleen, testes, and liver exhibited strong joining with the 10-nt substrate (Figure 7a), but as the microhomology length increased, we saw dramatic shifts in repair efficiency. Surprisingly, the spleen displayed a marked decline in joining efficiency with increasing microhomology length, indicating a clear preference for shorter homologies (see lane 7 in Figures 7a and 7 b-d).

**Fig. 7.**
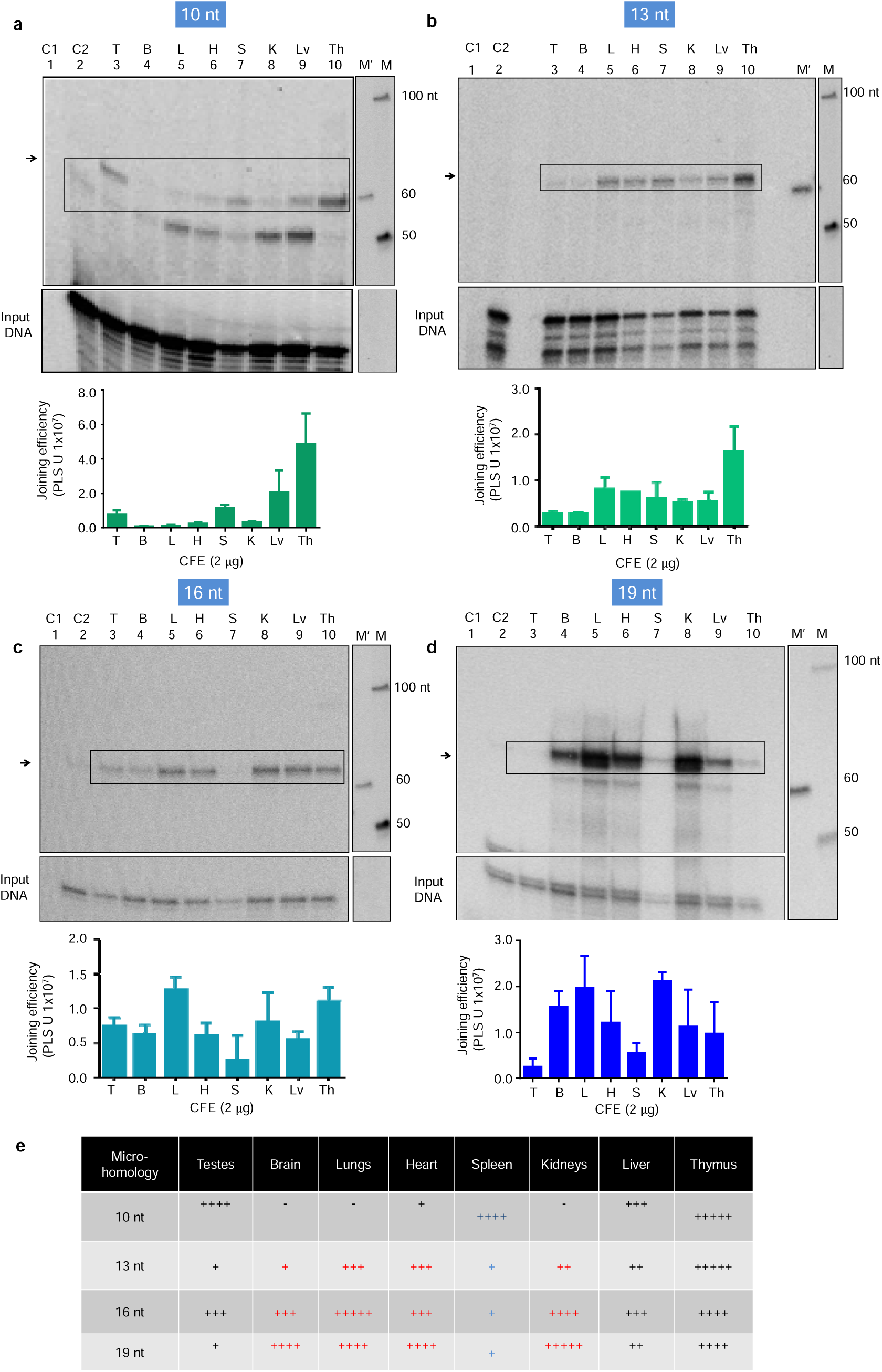
Comparison of MMEJ activity catalyzed by cell-free extracts from various rat organs using different microhomology substrates: **(a)** Comparison of MMEJ catalyzed by cell-free extracts from various rat organs using a 10-nt microhomology substrate. PCR amplification of the joint products gives a 62 nt joined product, marked with an arrow and a box. **(b)** Comparison of MMEJ catalyzed by CFE from various organs using a 13-nt microhomology substrate for 1 h at 30°C for testes and 37°C for other organs. The joined products (65 nt) were amplified by radioactive PCR. **(c)** Comparison of MMEJ catalyzed by CFE from various organs using a 16-nt microhomology substrate. The joined products (68 nt) were amplified by radioactive PCR. An arrow and a dotted box indicate MMEJ products. **(d)** Comparison of MMEJ catalyzed by CFE from various organs using a 19-nt microhomology substrate for 1 h at 30°C for testes and 37°C for other organs. The joined products (71 nt) were amplified by radioactive PCR. An arrow and a dotted box indicate MMEJ products. **(a-d)**. A bar graph quantifies results from at least three biological experiments, with joining efficiency labeled in PSLU. Error bars represent the mean ± SEM. The tissues analyzed in all panels are the brain (B), testis (T), thymus (Th), spleen (S), lungs (L), heart (H), liver (Lv), and kidney (K). M is a 50 bp ladder, and M’ is a 60 bp marker corresponding to the MMEJ products**. (e)** A table showing the comparison of MMEJ efficiency catalyzed by CFE from various organs using increasing microhomology regions. The number of plus represents the activity profile (five plus indicating maximum activity and one plus indicating minimum activity, whereas a minus indicates minimum/no joining. Red indicates an increase and blue indicates a decrease in joining efficiency.

In contrast, tissues like the lungs, kidneys, brain, and heart previously showing minimal repair with the 10-nt substrate (Figure 7a) demonstrated a substantial increase in repair efficiency with longer microhomologies, peaking at 16 or 19 nt (Figures 7c and 7d). Interestingly, the lung exhibited minimal repair with the 10-nt substrate but reached peak efficiency with the 16-nt substrate (compare lane 5 in Figures 7a and 7c). Similarly, the kidney displayed maximal repair activity with the 19-nt substrate (Figure 7d), and the brain and heart showed similar trends. These observations reveal a inverse relationship in repair efficiency between the 10-nt and 19-nt microhomology substrates across tissues: those favoring short microhomologies (10 nt) (e.g., testes, spleen) were less efficient at repairing with longer microhomologies, whereas those favoring longer microhomologies (e.g., lung, brain, kidney, heart) showed minimal activity at 10 nt (Figure 7d & 7a). Interestingly, testes showed a distinct non-linear pattern: robust joining at 10 nt, a sharp decline at 13 nt, partial recovery at 16 nt, and a secondary drop at 19 nt, suggesting a tightly regulated and length-sensitive repair mechanism (compare lane 3 in Figures 7a–d). In stark contrast, the thymus and liver maintained consistently high levels of MMEJ activity across all microhomology lengths (lanes 9 and 10 in Figures 7a–d), implying that repair efficiency in these tissues is mainly independent of microhomology length.

Importantly, this inverse trend in repair efficiency between short and long microhomologies appears to align with cellular context. Proliferative tissues such as the, testes and spleen favored shorter microhomologies (10 nt), while non-dividing or post-mitotic tissues, including the brain, heart, and kidney, preferentially utilized longer microhomologies (16&19 nt). These findings point to a regulatory shift in repair pathway usage that may be driven by differences in cell cycle status or chromatin context, further underscoring the dynamic nature of MMEJ regulation across tissues.

### Expression of DNA Repair Factors Correlates with tissue-specific preferences of microhomology

To investigate the molecular determinants underlying tissue-specific preferences in MMEJ, the expression level of key DNA repair proteins were analyzed across multiple murine tissues. These profiles were then correlated with end joining efficiency observed with defined substrates containing 10, 13, 16, or 19-nt microhomologies (Figure 8). Organs that demonstrated high MMEJ efficiency with short (10-nt) microhomologies, including the thymus, spleen, testes, and liver, exhibited elevated expression levels of essential MMEJ-associated factors. These included Ligase III, XRCC1, PARP1, MRE11, FEN1, Ligase I, and polymerase theta (Pol θ), which collectively participate in DNA end resection, gap filling, and ligation.

**Fig. 8.**
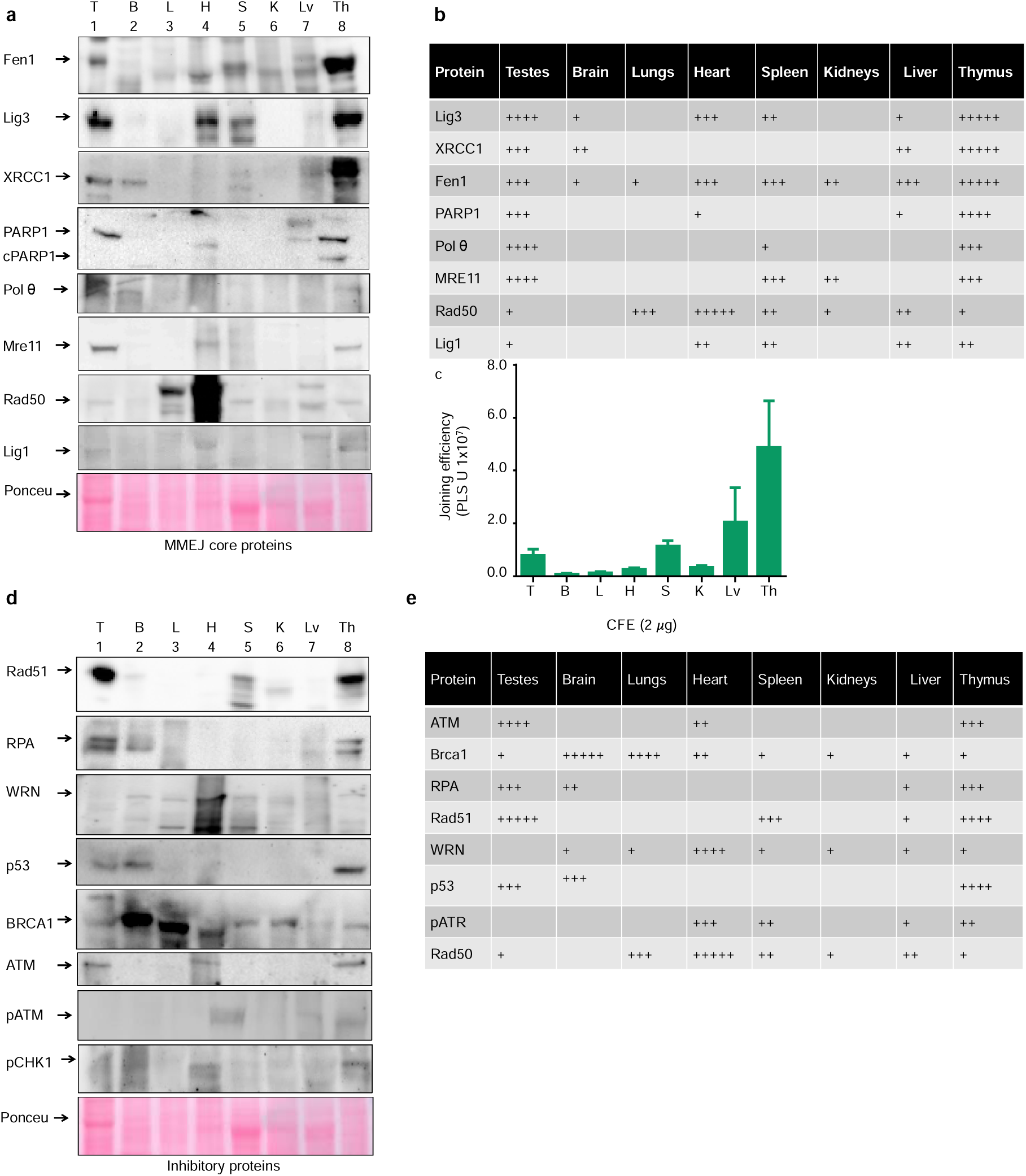
Western blot analysis of core MMEJ and inhibitory proteins across rat organs. **(a)** Western blots showing the expression of core MMEJ proteins in eight rat organs: testis (T), brain (B), lungs (L), heart (H), spleen (S), kidneys (K), liver (Lv), and thymus (Th). An arrow indicates the primary protein, while unmarked bands may represent isoforms. Ponceau staining is used as the loading control. **(b & c)** Correlation between MMEJ protein expression and repair efficiency profiles **(b)** Table showing the expression of core MMEJ proteins across organs, with plus signs (+) indicating relative protein abundance, with results from at least three independent experiments. **(c)** The lower panel depicts MMEJ activity profiles from each organ’s cell-free extracts (CFE). The Y axis is labeled with photostimulated luminescence units (PSLU), and error bars represent the standard error of the mean ± SEM, the results from at least three independent experiments. **(d)** Western blots showing the expression of MMEJ inhibitory proteins in the same organs, with the primary protein indicated by an arrow. Ponceau staining is the loading control. **(e)** The table shows the expression profile of MMEJ inhibitory proteins across organs, with plus signs (+) indicating relative protein abundance.

In contrast, the brain and heart expressed many of these same MMEJ components but failed to exhibit efficient repair with the 10-nt substrate. Closer examination revealed that PARP1 was absent in the brain and XRCC1 was lacking in the heart, suggesting that the absence of even a single core component may compromise MMEJ activity. These findings highlight the requirement for a complete element of MMEJ machinery for optimal repair efficiency.

### Repair Activity in Tissues with Limited Canonical MMEJ Machinery

Interestingly, robust repair of longer microhomology substrates (16–19 nt) was observed in the lungs and kidneys, despite the absence or low expression of several canonical MMEJ proteins. These results suggest that alternative, MMEJ-independent repair pathways may be engaged in these tissues without MMEJ machinery. This tissue-specific divergence in repair strategy underscores the flexibility of the cellular DNA repair network in adapting to local molecular environments.

### Influence of Inhibitory Factors on MMEJ Suppression

To explore potential regulators of tissue-specific MMEJ efficiency, we assessed the expression of key proteins inhibiting DNA end resection or promoting alternative DSB repair pathways such as HR and NHEJ. Strikingly, tissues with high MMEJ, including the thymus, spleen, and testes, also exhibited elevated levels of RPA, RAD51, ATM, WRN, and p53 (Figure 8a, b), suggesting that MMEJ operates in a cellular context enriched for components of major DSB repair pathways.

WRN, a helicase known to suppress MMEJ by blocking the recruitment of MRE11 and CtIP to DNA breaks and thereby inhibiting the resection required for MMEJ initiation [56], was notably up-regulated in the heart, a tissue with low MMEJ. This observation shows that WRN may actively suppress MMEJ in certain tissues, even with MMEJ-supporting factors. Together, these data suggest that MMEJ activity is shaped by the presence of core MMEJ machinery and the broader regulatory environment influencing DSB pathway choice.

### Role of BRCA1 in Regulating Microhomology Usage

Although BRCA1 is primarily recognized for its role in promoting HR, emerging evidence suggests it can suppress MMEJ through Chk2-dependent signaling pathways that limit microhomology usage at DNA double-strand breaks [57, 58]. In my analysis, tissues such as the lung and brain, which showed poor repair efficiency with 10-nucleotide microhomologies, exhibited high levels of BRCA1 expression. This inverse relationship suggests that BRCA1 may actively repress MMEJ in favor of more accurate repair mechanisms, particularly in post-mitotic or genomically sensitive tissues where the fidelity of repair is paramount. These findings support a context-dependent role for BRCA1 in modulating DSB repair pathway choice, potentially safeguarding genomic integrity by limiting reliance on error-prone repair routes such as MMEJ.

## Discussion

MMEJ has traditionally been viewed as a backup DSB repair pathway, predominantly active in cancer cells or under conditions where canonical mechanisms such as c-NHEJ or HR are compromised [25–27]. However, the growing evidence presented in this study offers a revised understanding and supports a different perspective [6, 30, 40–41, 60–62]. Despite this paradigm shift, the physiological contexts in which this inherently mutagenic pathway is activated remain incompletely understood.

Recent studies have proposed that MMEJ plays a critical role in repairing mitotic DSBs, which occur during periods of highly condensed chromatin, where accessibility to repair factors may be limited [30, 63]. The findings presented here support this hypothesis, revealing that MMEJ is markedly enriched in proliferative, mitotically active tissues such as the thymus, spleen, testes, and liver, while being substantially reduced or absent in largely post-mitotic tissues like the brain, heart, kidneys, and lungs (Figure 2). This pattern suggests that MMEJ may not be merely a compensatory mechanism but rather a regulated, context-specific strategy aligned with the proliferative status of the tissue. In such dividing environments, the mutagenic potential of MMEJ may be tolerated due to high cellular turnover, reducing the long-term impact of genomic errors.

In addition to revealing the tissue-specific regulation of MMEJ, this study uncovers key factors that may underlie this differential activity. The data show that the expression of MMEJ-associated proteins such as Ligase III, XRCC1, PARP1, MRE11, FEN1, Ligase I, and polymerase theta (Pol θ) and inhibitory proteins (RPA, RAD51, ATM, WRN, and p53 are closely linked to the variation in repair efficiency observed across tissues (Figure 8).

The length of microhomology at DSB ends is responsible for the tissue-specific function of MMEJ. MMEJ activity was observed at microhomology lengths of approximately 10 nucleotides. Intriguingly, post-mitotic tissues exhibited low MMEJ at 10 ntmicrohomology length but showed increased repair with longer microhomologies (Figure 9). This indicates that microhomology length preference is tissue-specific and possibly governed by distinct regulatory mechanisms. Supporting this, we observed Pol θ, known for aligning short microhomologies [55], is highly expressed in proliferative tissues like the thymus and testes, but is largely absent from most non-dividing tissues (Figure 8a). Conversely, the brain, lungs, and heart show elevated levels of BRCA1 (Figure 8d), which is known to suppress microhomology usage, possibly favouring high alternative repair in long-lived, non-dividing cells. These observations point to a highly coordinated regulatory framework that limits mutagenic repair where genome stability is paramount.

**Fig. 9.**
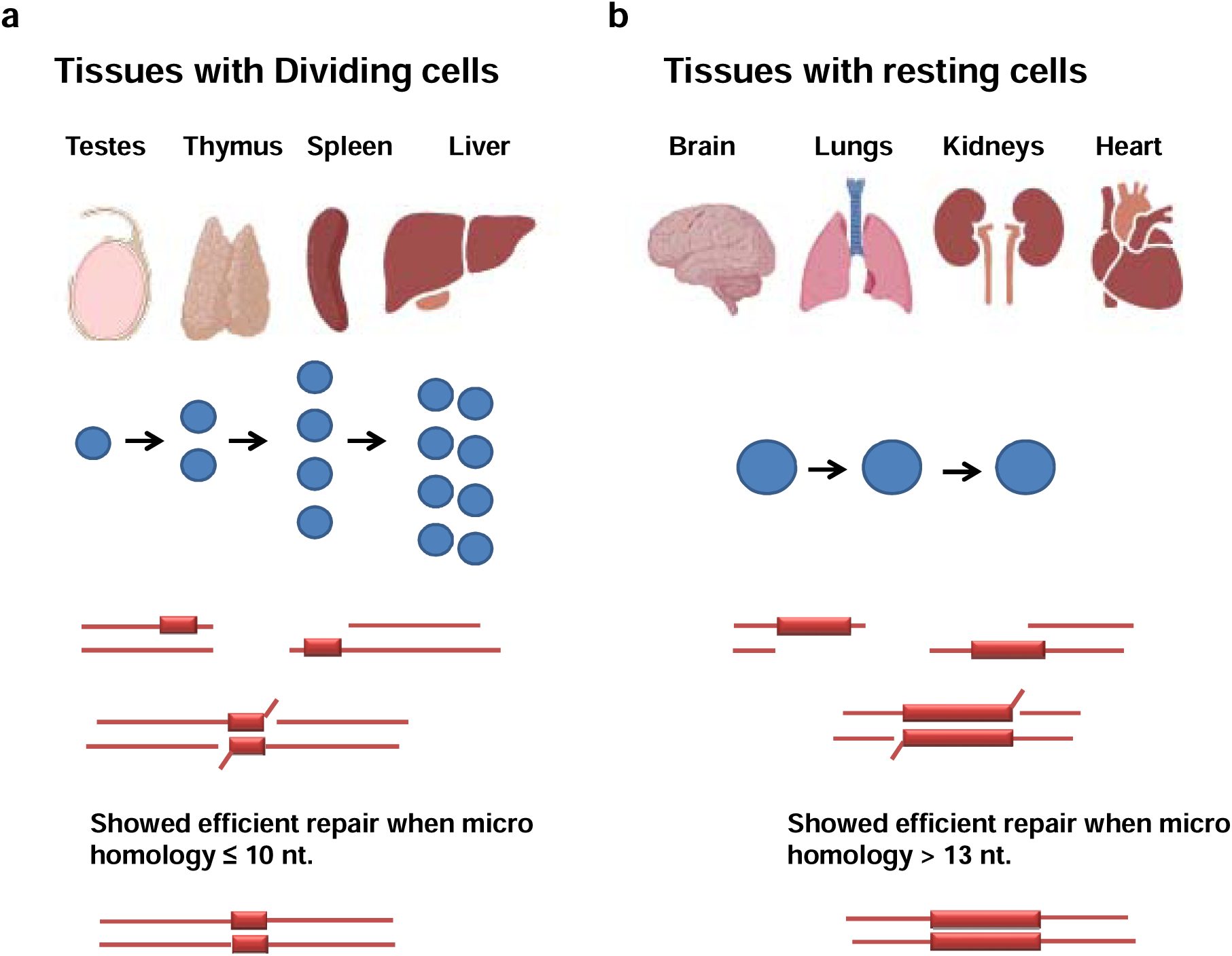
Summary schematic of tissue-specific regulation of the MMEJ pathway in normal rat tissues. This Fig. 1. illustrates the study’s core finding: MMEJ is differentially regulated across tissues based on their proliferative state. (a) Proliferative tissues such as the thymus, testes, spleen, and liver showed high MMEJ activity, efficiently repairing DNA substrates with short 10-nucleotide microhomologies. This correlates with elevated expression of key MMEJ proteins in these tissues. (b) In contrast, non-proliferative tissues, including brain, heart, lungs, and kidneys, exhibited poor repair at 10-nt microhomologies, but repair efficiency significantly improved with longer microhomology lengths (≥13 nt). These results suggest that MMEJ pathway activity and microhomology length preference are tightly linked to tissue proliferation status, with potential implications for tissue-specific genome maintenance.

Interestingly, tissues such as the lungs and kidneys, which express low levels of canonical MMEJ components, exhibit elevated end joining when longer microhomologies are present (Figure 7c & 7d). This observation suggests the possible involvement of alternative repair pathways, such as single-strand annealing (SSA). Rad52, a key mediator of SSA, is known to facilitate the alignment of extended microhomology tracts [48]. It may be more active or highly expressed in post-mitotic tissues. These findings point to an underexplored dimension of DSB repair that extends beyond the conventional MMEJ framework and highlight the potential for tissue-specific reliance on distinct repair mechanisms.

Microhomology length is a critical determinant not only of MMEJ efficiency but also of broader repair pathway choice. While previous studies suggested that MMEJ efficiency generally increases with longer microhomologies [47], my data reveal exceptions. For example, MMEJ efficiency decreases in the spleen as microhomology length increases beyond 10 nt (Compare Lane 7 of 7a with 7b, c, and d). This may be attributed to the absence of KU80 and XRCC4 in splenic B cells [63]. In the absence of these proteins, antibody class switch recombination (CSR) preferentially utilizes shorter microhomologies [64].

Adding complexity, Rad52, a protein central to SSA and HR, is also required for CSR, a process commonly associated with MMEJ [64], [65]. This overlap challenges the binary distinction between MMEJ and SSA, raising the possibility that these pathways are interlinked or co-regulated depending on physiological needs. Microhomology usage may thus serve as a key determinant in the dynamic selection of repair pathways.

The testes present another unique exception: I observed a non-linear relationship between microhomology length and repair efficiency. End joining efficiency peaked at 10 nt, dipped at 13 nt, rose again at 16 nt, and declined at 19 nt (Figure 7). Upon further analysis, it was revealed that specific sequence characteristics influence the observed non-linearity of MMEJ activity with changing microhomology length. Notably, the 13-nt substrate, which was GC-rich, exhibited reduced MMEJ efficiency. These findings contrast with previous studies that have associated higher GC content with enhanced MMEJ activity [67]. These results suggest that, in the testes, MMEJ may be less efficient at processing GC-rich microhomologies, indicating a sequence-dependent constraint on repair efficiency. Additionally, the decline in efficiency at 19 nt may reflect the challenges associated with processing longer microhomologies, as seen with the 19-nt substrates, which further impairs repair in germ cells. Given the heritability of germ-line mutations, tight regulation of MMEJ in the testes is likely essential for preserving genome fidelity.

Beyond its role in mitosis, MMEJ also contributes to several physiological processes, which likely explain its tissue-specific enrichment in this study. The high MMEJ activity detected in the liver is due to its regenerative capacity and rapid cell turnover. MMEJ is implicated in meiotic DSB repair in the testes, supported by its co-localization with SPO11-induced breaks and high Pol θ expression [68], [42]. Furthermore, the presence of MMEJ activity in both sperm and *Xenopus* egg extracts suggests an evolutionarily conserved role for this pathway in germline genome maintenance [41].

MMEJ facilitates class switch recombination (CSR) in B cells within the spleen. This study demonstrated robust MMEJ activity alongside reduced levels of Pol θ (Figure 8a). These findings suggest that alternative polymerases, such as Pol β, δ, or λ, may compensate for Pol θ in mediating end joining [11]–[13]. In the thymus, while V(D)J recombination primarily relies on c-NHEJ [69], [70], substantial MMEJ was observed, likely originating from non-lymphoid cells such as thymic epithelial cells (TECs). This suggests that while c-NHEJ predominates in lymphoid cells, MMEJ may contribute to the repair processes in other cell types within the thymus [71].

The study suggests that MMEJ activity is determined by the interplay between local sequence context and protein availability. It also highlights tissue-specific differences in the expression of MMEJ-associated proteins. However, the regulatory mechanisms governing microhomology length in mitotic versus post-mitotic tissues remain inadequately defined. In particular, the factors that dictate the preferential formation of DSBs flanked by short microhomology in mitotic tissues, as opposed to longer microhomology in post-mitotic tissues, are poorly understood. This raises important questions about the precise molecular processes that regulate microhomology availability and how these mechanisms are spatially and temporally controlled across different cells. This indicates that MMEJ might be involved in repairing DSBs that are not random events or merely a result of external DNA damage but are likely tightly regulated, potentially through enzymatic processes like DSB formation in CSR.

A compelling area for future investigation is deciphering how and why MMEJ functions during mitosis when chromatin is highly compacted and DNA access is limited. Studies in compact-genome organisms like *Oikopleura dioica* show a reliance on MMEJ over c-NHEJ [72], reinforcing that chromatin structure directly influences repair pathway choice. Mitotic condensation may expose fragile genomic regions, increasing susceptibility to DSBs and necessitating repair via pathways like MMEJ.

Although the current study provides critical insights into regulating MMEJ in normal tissues, it relies primarily on in vitro models. Future in vivo experiments, akin to those by Vidya et al. [73], will be essential to validate the current findings. To date, no study has definitively demonstrated SSA activity in healthy tissues, and further research is needed to better understand the physiological relationship between MMEJ and SSA.

This study redefines Microhomology-Mediated End Joining (MMEJ) as a tightly regulated, context-dependent DNA repair pathway with physiological relevance beyond its canonical role in cancer. We show that MMEJ is preferentially active in proliferative tissues—such as thymus, spleen, testes, and liver—where its mutagenic potential is tolerated due to high cellular turnover. In contrast, post-mitotic tissues like brain, heart, lungs, and kidneys suppress MMEJ through elevated expression of inhibitory factors such as WRN and BRCA1. We demonstrate that microhomology length and sequence context critically influence pathway selection, with longer microhomologies favoring SSA, especially in non-dividing tissues. Our findings also reveal non-linear and tissue-specific patterns of MMEJ efficiency, suggesting that repair pathway choice is shaped by both chromatin context and the availability of core and regulatory proteins. Collectively, this work provides new insights into the dynamic landscape of DSB repair in normal tissues and highlights MMEJ as a physiologically significant but carefully restrained repair strategy. These insights have broad implications for genome maintenance, tissue regeneration, and the development of precision therapies targeting repair pathways in both proliferative and post-mitotic settings.

## Abbreviations

DSB: Double-Strand Break
HR: Homologous Recombination
c-NHEJ / NHEJ: (Canonical) Non-Homologous End Joining
MMEJ: Microhomology-Mediated End Joining
SSA: Single-Strand Annealing
Pol θ: DNA Polymerase Theta
CFE: Cell-Free Extract
TECs: Thymic Epithelial Cells
CSR: Class Switch Recombination
GC: Guanine-Cytosine (content)
ATM: Ataxia Telangiectasia Mutated
RPA: Replication Protein A
BRCA1: Breast Cancer Gene 1
XRCC1: X-Ray Repair Cross Complementing 1
FEN1: Flap Endonuclease 1
RAD51: Radiation Sensitive 51
KU80: Ku Autoantigen, 80 kDa
MRN complex: MRE11-RAD50-NBS1 Complex
CtIP: C-terminal Binding Protein (CtBP)-interacting Protein
PARP1: Poly (ADP-ribose) Polymerase 1
RAG: Recombination Activating Gene
SPO11: Meiotic recombination protein
nt: Nucleotide(s)

## Acknowledgements

I would like to thank Prof. Sathees C. Raghavan for his invaluable, suggestions, discussions and critical input during the course of this work. I am also grateful to him for providing essential reagents and laboratory space to carry out the experiments. This work was supported by the Council of Scientific and Industrial Research (CSIR), the Grant-in-Aid for Research Program (GARP), the Department of Biotechnology (DBT), and the Indian Institute of Science (IISc), India.

---

**Supplementary fig. 1.**
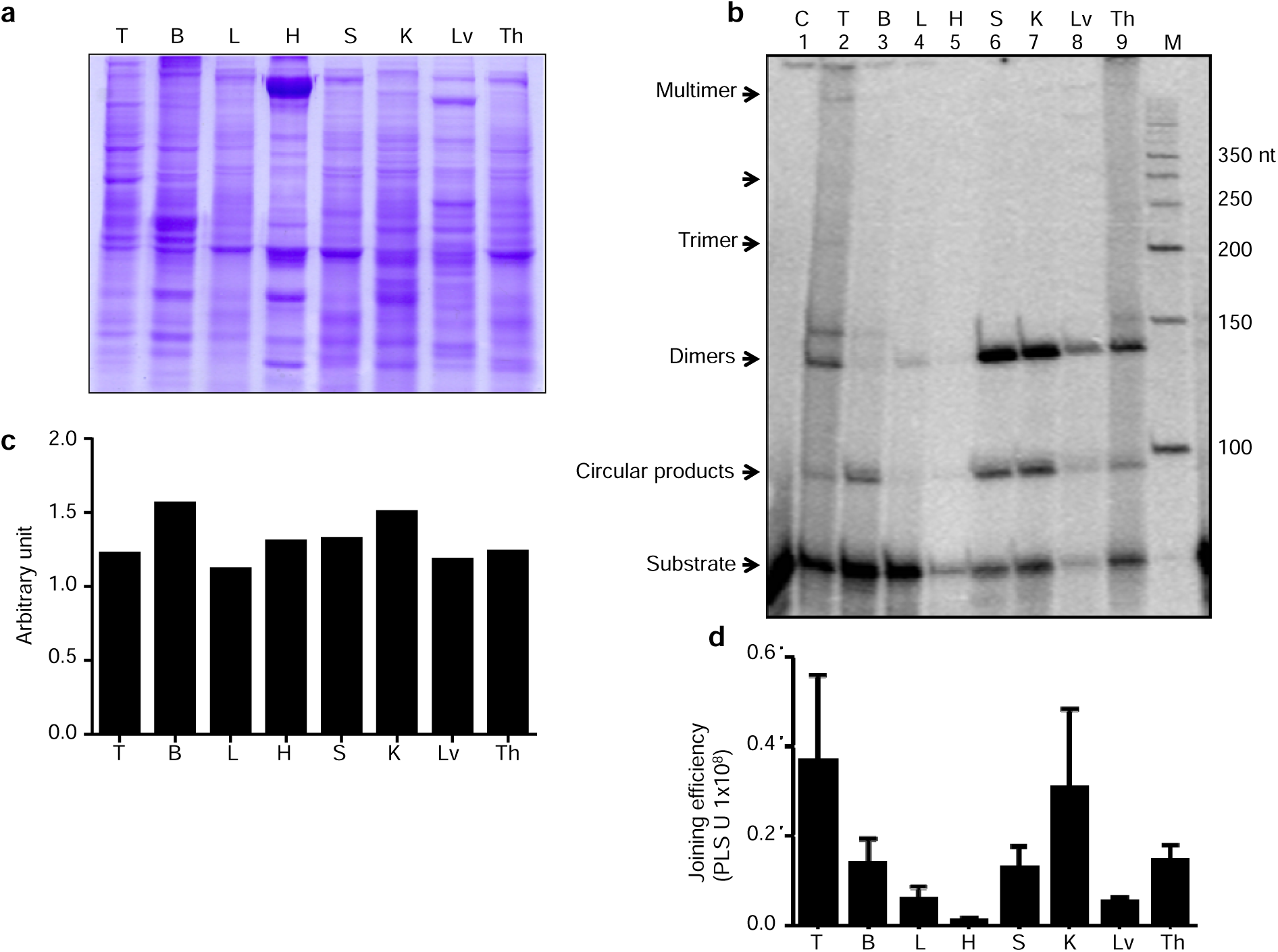
NHEJ assay to confirm the function activity of proteins in CFE: (**a**) SDS-PAGE analysis demonstrates the normalization of CFE from various organs rats, including testes(T), brains (B), lungs (L), heart (H), spleen (S), kidneys (K), liver (Lv), and thymus (Th). (**b**) Representative denaturing PAGE shows NHEJ efficiency after incubating 2 µg CFE from different rat organs with a dsDNA substrate containing compatible ends. Lane 1 represents the no protein control, and M indicates the 50 nt marker. (**c**) A bar graph illustrates the protein normalization profile obtained from the SDS-PAGE analysis shown in panel (**a**). (**d**) A bar graph showing the quantification of NHEJ joining products (measured in photostimulated luminescence units, PSLU) after incubating a dsDNA substrate with compatible ends and normalized CFE from various early-age rat organs. The experiment was repeated three times using three different batches of cell-free extracts, and error bars represent the mean ± SEM.

**Supplementary fig. 2.**
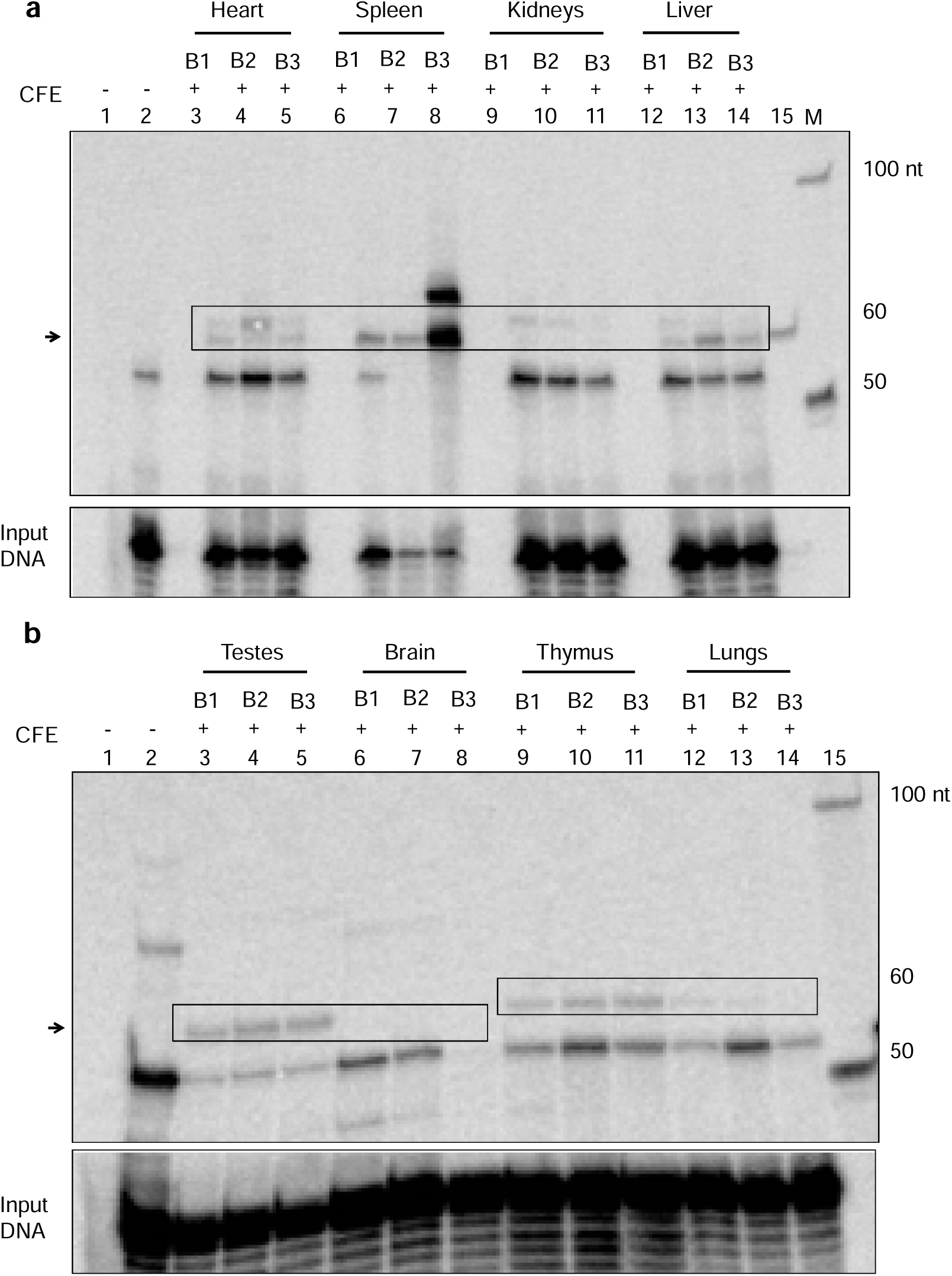
Comparison of MMEJ activity between the batches of CFE: Three independent batches of cell-free extract (CFE) were prepared from each of the following organs: testes, brain, lungs, heart, spleen, kidneys, liver, and thymus. (a) MMEJ activity is compared between the batches of CFE from heart (lanes 3–5), spleen (lanes 6–8), kidneys (lanes 9–11), and liver (lanes 12–15). Lane 1 represents the no-template control, and lane 2 represents the no-protein control. (b) MMEJ activity is compared between the batches of CFE from testes (lanes 3–5), brain (lanes 6–8), thymus (lanes 9–11), and lungs (lanes 12–15). Lane 1 is the no-template control, and lane 2 is the no-protein control. B1, B2, and B3 correspond to batches 1, 2, and 3 of CFE from each tissue, respectively.

