## supplementary information for "Context-Dependent Regulation of Microhomology-Mediated End Joining in Normal Tissues: Insights into Tissue-Specific Activation of DNA Repair Pathways"

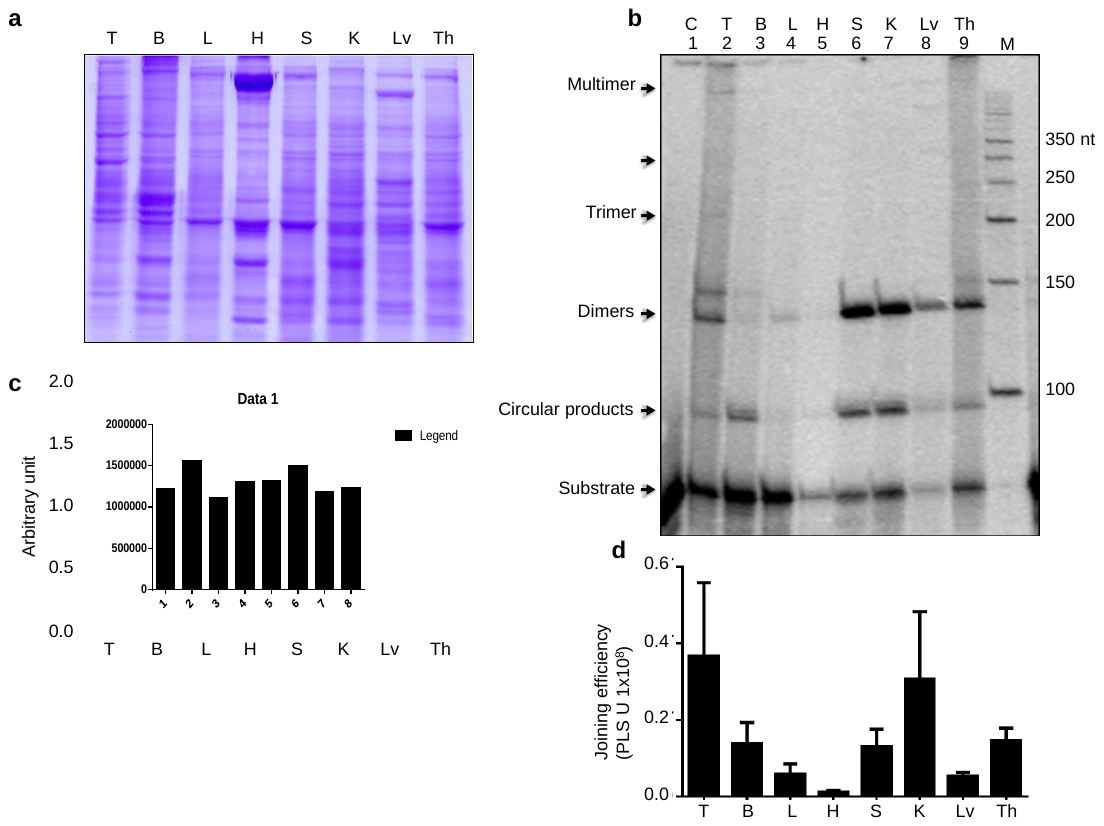

**Supplementary figure1:NHEJ assay to confirm the function activity of proteins in CFE:** (**a**) SDS-PAGE analysis demonstrates the normalization of CFE from various organs rats, including testes(T), brains (B), lungs (L), heart (H), spleen (S), kidneys (K), liver (Lv), and thymus (Th). (**b**) Representative denaturing PAGE shows NHEJ efficiency after incubating 2 µg CFE from different rat organs with a dsDNA substrate containing compatible ends. Lane 1 represents the no protein control, and M indicates the 50 nt marker. (**c**) A bar graph illustrates the protein normalization profile obtained from the SDS-PAGE analysis shown in panel (**a**). (**d**) A bar graph showing the quantification of NHEJ joining products (measured in photostimulated luminescence units, PSLU) after incubating a dsDNA substrate with compatible ends and normalized CFE from various early-age rat organs. The experiment was repeated three times using three different batches of cell-free extracts, and error bars represent the mean ± SEM.

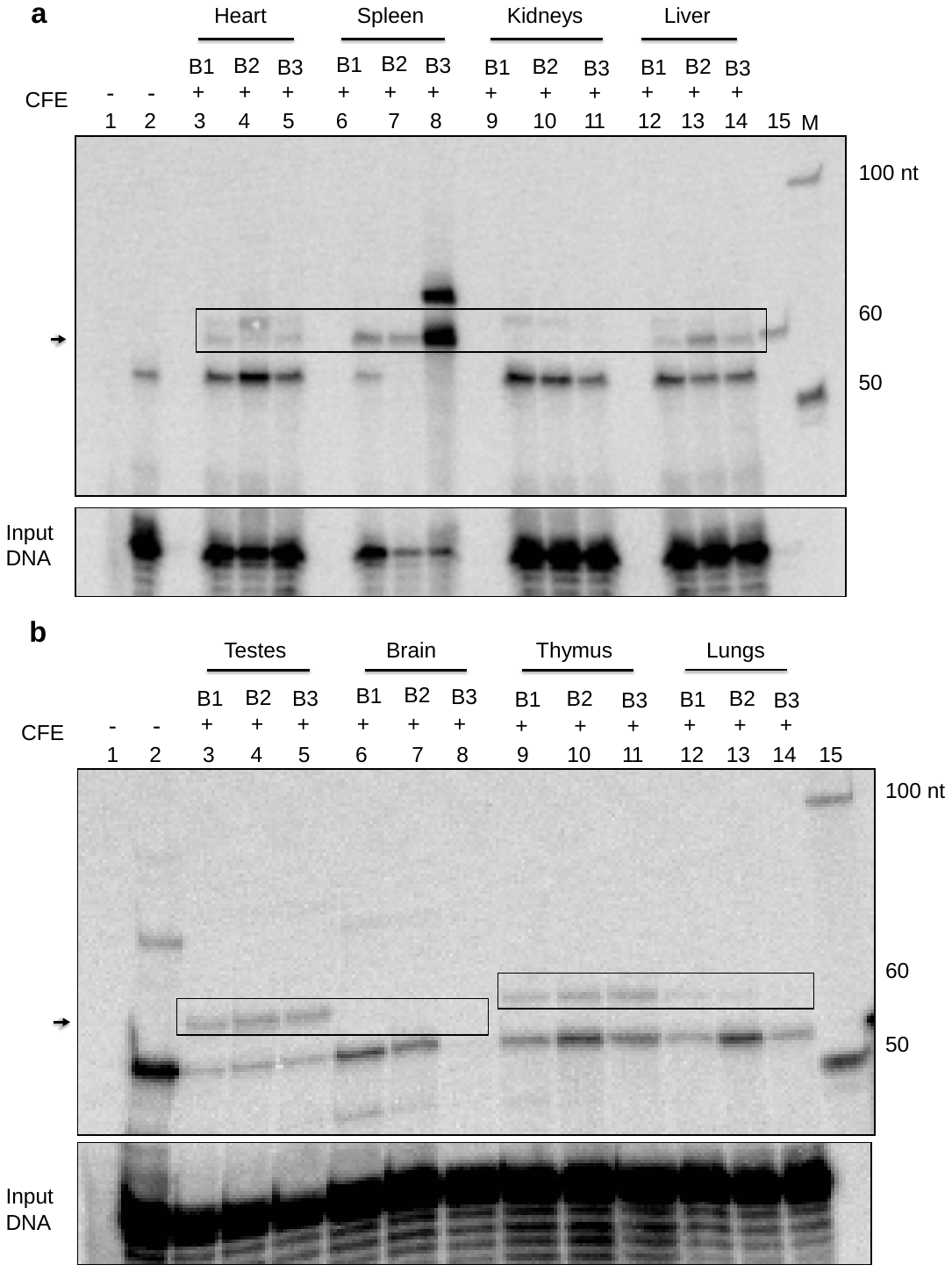

**Supplementary figure2: Comparison of MMEJ activity between the batches of CFE:** Three independent batches of cell-free extract (CFE) were prepared from each of the following organs: testes, brain, lungs, heart, spleen, kidneys, liver, and thymus. (a) MMEJ activity is compared between the batches of CFE from heart (lanes 3–5), spleen (lanes 6–8), kidneys (lanes 9–11), and liver (lanes 12–15). Lane 1 represents the no-template control, and lane 2 represents the no-protein control. (b) MMEJ activity is compared between the batches of CFE from testes (lanes 3–5), brain (lanes 6–8), thymus (lanes 9–11), and lungs (lanes 12–15). Lane 1 is the no-template control, and lane 2 is the no-protein control. B1, B2, and B3 correspond to batches 1, 2, and 3 of CFE from each tissue, respectively.

**Table1**

| Name | Sequence |
| --- | --- |
| SS54 | 5' –GATTAGATAATCAGTTCATCGAGTCCTCACGTAGATCGGTACTACTCGAGCTGAG-3’ |
| SS60 | 5’ -AACTGCATCTGTAGGTCG-3’ |
| SS61 | 5' -GAGTAGTACCGATCTACGTG-3’ |
| SS62 | 5' -TTAACTCAGCTCGAGTAGTACCGATCTACGTGAGGACTCGATGAA CTGATTATCT3’ |
| SS65 | 5’ -AGCTAACTGCATCTGTAGGTCGCGAATCAGTTCACTGCTTAA-3’ |
| SS66 | 5’ -GCAGTGAACTGATTCGCGACCTACAGATGCAGTT-3’ |
| SS69 | 5’ -AGCTAACTGCATCTGTAGGTCGCCCAGATCTGCAGTGATGCTTAA-3 |
| SS70 | 5' -GCATCACTGCAGATCTGGGCGACCTACAGATGCAGTT-3' |
| SS71 | 5' GATTAGATCAGATCTGCAGTGGCGAGTCCTCACGTAGATCGGTACTACTCGAGCTGAG-3’ |
| SS72 | 5' -GCTCGAGTAGTACCGATCTACGTGAGGACTCGCCACTGCAGATCTGATCT-3' |
| SS73 | 5’ -AGCTAACTGCATCTGTAGGTCGCTAGGGATAACAGGGTAAGTGCTTAA-3’ |
| SS74 | 5' -GCACTTACCCTGTTATCCCTAGCGACCTACAGATGCAGTT-3‘ |
| SS75 | 5' -GATTAGACAGGGATAACAGGGTAATCGAGTCCTCACGTAGATCGGT  ACTACTCGAGCTGAG-3' |
| SS76 | 5' -GCTCGAGTAGTACCGATCTACGTGAGGACTCGATTACCCTGTTATCCCTGTCT-3’ |
| SS92 | 5' -AGCTAACTGCATCTGTAGGTCGCTAACTATAACGGTCCTAAGGTAGCCTCATTAA-3’ |
| SS93 | 5' -TGAGGCTACCTTAGGACCGTTATAGTTAGCGACCTACAGATGCAGTT-3' |
| SS94 | 5'AGCTAGAGATAACGGTCCTAAGGTAGCGAGAGTCCTCACGTAGATCGGTACTACTCGAGCTGAG-3’ |
| SS95 | 5'-GCTCGAGTAGTACCGATCTACGTGAGGACTCTCGCTACCTTAGGACCGTTATCTCT-3’ |
| SS96 | 5'AGCTAGAGACTATAACGGTCCTAAGGTAGCGAGAGTCCTCACGTAGATCGGTACTACTCGAGCTGAG-3’ |
| SS97 | 5'GCTCGAGTAGTACCGATCTACGTGAGGACTCTCGCTACCTTAGGACCGTTATAGTCTCT-3’ |
| SCR 19 | 5’-GATCCCTCTAGATATCGGGCCCTCGATCCGGTACTACTCGAGCCGGCTAGCTTCGAT- GCTGCAGTCTAGCCTGAG-3' |
| SCR 20 | 5’-GATCCTCAGGCTAGACTGCAGCATCGAAGCTAGCCGGCTCGAGTAGTACCGGATCGAGGGCCCGATAT CTAGAGG-3’ |

|  | 1. |
| --- | --- |
